# Altered iron-sulfur cluster transfer in Arabidopsis mitochondria reveals lipoyl synthase as a Janus-faced enzyme that generates toxic sulfide

**DOI:** 10.1101/2023.08.30.555573

**Authors:** Luca Pedroletti, Anna Moseler, Stefan Timm, Gernot Poschet, Maria Homagk, Jeremy X. L. The, Stephan Wagner, Markus Wirtz, Rüdiger Hell, Andreas J. Meyer

## Abstract

Iron–sulfur (Fe–S) cluster are vital cofactors in all domains of life. Mitochondrial Fe–S cluster assembly occurs in two major steps to first build [2Fe–2S] clusters and subsequently assemble these into [4Fe–4S] clusters. The two assembly machineries are interconnected by glutaredoxin S15 (GRXS15) that transfers [2Fe–2S] clusters to the second machinery. Diminished cluster transfer activity of GRXS15 in Arabidopsis mitochondria causes specific defects associated with lipoyl synthase (LIP1) activity. Conversely, overexpression of *LIP1* in wild-type plants causes the release of toxic amounts of sulfide that can be detoxified by increasing the capacity for sulfide fixation through overexpression of *O*-acetylserine-(thiol)-lyase. The release of sulfide by lipoyl synthase causes a disturbance of mitochondrial sulfide homeostasis resulting in distinct and readily observable macroscopic phenotypes. These phenotypes enable a direct readout of consequences resulting from defects in Fe–S cluster assembly or targeted modulation of Fe–S cluster flux in mitochondria.

## Introduction

Multiple essential mitochondrial proteins depend on iron–sulfur (Fe–S) clusters as cofactors to enable electron transfer reactions or catalytic activities (Przybyla-Toscano et al., 2020). Assembly of Fe–S clusters occurs in two major steps with the initial synthesis of [2Fe–2S] clusters by a first machinery followed by a subsequent assembly into [4Fe–4S] clusters by a second machinery (Balk and Schaedler, 2014; Pedroletti et al., 2023). Glutaredoxin S15 (GRXS15) has been identified as an essential transfer chaperone for [2Fe–2S] clusters to the [4Fe–4S] cluster assembly machinery in *Arabidopsis thaliana* (Arabidopsis) (Moseler et al., 2015; Azam et al., 2020; Trnka et al., 2020). Complementation of embryo-lethal null mutants with weak mutant alleles and severe knockdown of *GRXS15* both lead to dwarfism and defects in mitochondrial protein lipoylation (Moseler et al., 2015; Ströher et al., 2016; Moseler et al., 2021). Similarly, mutants with severely diminished levels of the [4Fe–4S] cluster carriers NifU-like4 (NFU4) and NifU-like5 (NFU5) cause an impairment of mitochondrial glycine decarboxylase (GDC) activity, which also points at dysfunction of lipoyl synthase (LIP1) (Przybyla-Toscano et al., 2022). While diminished LIP1 activity in *nfu* mutants may be explained with the specificity of NFUs for cluster transfer to LIP1 (Przybyla-Toscano et al., 2022), it is less obvious why a bottleneck upstream of the [4Fe–4S] assembly machinery at GRXS15 should have similar specific metabolic consequences in protein lipoylation rather than broader effects at multiple mitochondrial Fe–S proteins.

Protein lipoylation in mitochondria occurs on subunits of four multiprotein complexes, including GDC, pyruvate dehydrogenase (PDC), 2-oxoglutarate dehydrogenase (OGDC), and branched-chain α-keto acid dehydrogenase (BCKDC) (Douce et al., 2001; Solmonson and DeBerardinis, 2018). Defects in protein lipoylation consequently affect all four complexes and result in a characteristic alteration of metabolic signatures in the respective mutants (Fu et al., 2020; Lill and Freibert, 2020; Moseler et al., 2021; Przybyla-Toscano et al., 2022). The highly conserved LIP1 belongs to the family of radical *S*-adenosylmethionine (SAM) enzymes, which all contain a [4Fe–4S] cluster and catalyse formation of a 5′-deoxyadenosyl radical to target largely inert substrates for hydrogen abstraction. LIP1 carries an additional ‘auxiliary’ [4Fe–4S] cluster that provides the two sulfur atoms required for synthesis of lipoic acid (LA) (McLaughlin et al., 2016; McCarthy and Booker, 2017). In mitochondria of Arabidopsis suspension culture cells, LIP1 is very lowly abundant with only 85 copies per mitochondrion (Fuchs et al., 2020). Nevertheless, it has been proposed that LIP1 may be a major sink for [4Fe–4S] clusters if one cluster is sacrificed in each catalytic cycle (Pedroletti et al., 2023). The fate of the fragmentary cluster after abstraction of two sulfur atoms is unknown and it possibly falls apart, releasing ferrous iron and sulfide.

Free sulfide, which under alkaline conditions is predominantly present as hydrosulfide anion (HS^−^), inhibits complex IV (cytochrome *c* oxidase, COX) of the mitochondrial electron transport chain (mETC) with IC_50_ values between 6.9 nM and 200 nM (Kabil and Banerjee, 2010; Birke et al., 2012; Nicholls et al., 2013; Domán et al., 2023). To prevent inhibition of COX and a block in ATP synthesis, mitochondria require appropriate mechanisms for removal of sulfide. In fission yeast (*Schizosaccharomyces pombe*) and human mitochondria, sulfide is detoxified via sulfide quinone oxidoreductase (SQR), which channels electrons from sulfide directly into the mETC and transfers the resulting sulfane (S^0^) sulfur to reduced glutathione (GSH) to generate *S*-sulfanylglutathione (Theissen et al., 2003; Vitvitsky et al., 2021; Zhang et al., 2021). Plants do not have SQR but contain a mitochondrial cysteine synthase complex (CSC) consisting of serine acetyltransferase2;2 (SERAT2;2) and *O*-acetylserine (thiol) lyase C (OAS-TL C) to fix sulfide (Heeg et al., 2008; Watanabe et al., 2008). Because cysteine biosynthesis in plants is assumed to be largely confined to chloroplasts and cytosol rather than mitochondria (Krüger et al., 2009; Takahashi et al., 2011) and because OAS-TL C in mitochondria accounts for only 5 % of the total OAS-TL activity, the role for mitochondrial cysteine biosynthesis has remained enigmatic (Birke et al., 2012).

Here, we investigated the distribution of [4Fe–4S] clusters between different apoproteins in plant mitochondria and the question why diminished transfer of [2Fe–2S] clusters by GRXS15 specifically results in deficient protein lipoylation. We hypothesized that LIP1, despite its very low abundance, is effectively a large sink for [4Fe–4S] clusters, because it sacrifices a cluster in each catalytic cycle. We further proposed that sulfide release from cluster disintegration creates the primary demand for cysteine biosynthesis in mitochondria. To test these hypotheses, we exploited different mutants affected in GRXS15-mediated [2Fe–2S] transfer, altered the [4Fe–4S] distribution by modifying the abundance of different apoproteins, and investigated the respective metabolic effects. The model for sulfide release by LIP1 and the essential role of cysteine biosynthesis in the matrix was validated genetically and by physiological characterization of the engineered plants.

## Results

### Diminished capacity of GRXS15 for iron–sulfur cluster transfer causes a partial photorespiratory phenotype

To dissect the consequence of GRXS15 deficiencies, we first compiled a collection of *grxs15* mutants with different abundance and activity of GRXS15 that had been described earlier (Moseler et al., 2015; Ströher et al., 2016; Moseler et al., 2021). The T-DNA insertion mutant *grxs15-1* and the artificial microRNA interference mutant *amiR* are knockdown lines with residual GRXS15 while *K83A #3* and *#4* are *grxs15* null mutants that were complemented with a GRXS15 variant (K83A) compromised in its ability to coordinate the [2Fe–2S] cluster. Severity of the phenotype in *K83A #3* and *#4* inversely correlates with the expression levels of the complementation construct, which is highest in the least severe mutant *#3* (Moseler et al., 2015). Generally, the complementation lines *K83A #3* and *#4* are more severely affected than the knockdown lines (Supplemental Figure S1).

In *amiR*, *K83A #3* and *#4*, compromised lipoylation of mitochondrial proteins results in accumulation of substrates of the four lipoyl-dependent dehydrogenase enzyme complexes, pyruvate, 2-oxoglutarate (2-OG), glycine and branched-chain α-keto acids (BCKAs) (Ströher et al., 2016; Moseler et al., 2021). Because glycine decarboxylation by GDC is an essential step in photorespiration (Douce et al., 2001; Bauwe, 2023) (Figure 1A), we reasoned that the severe dwarf phenotype of *grxs15* mutants may be interpreted, at least to some extent, as a photorespiratory phenotype. To test this hypothesis, we initially grew the *grxs15* mutants under elevated CO_2_ to suppress photorespiration.

**Figure 1.**
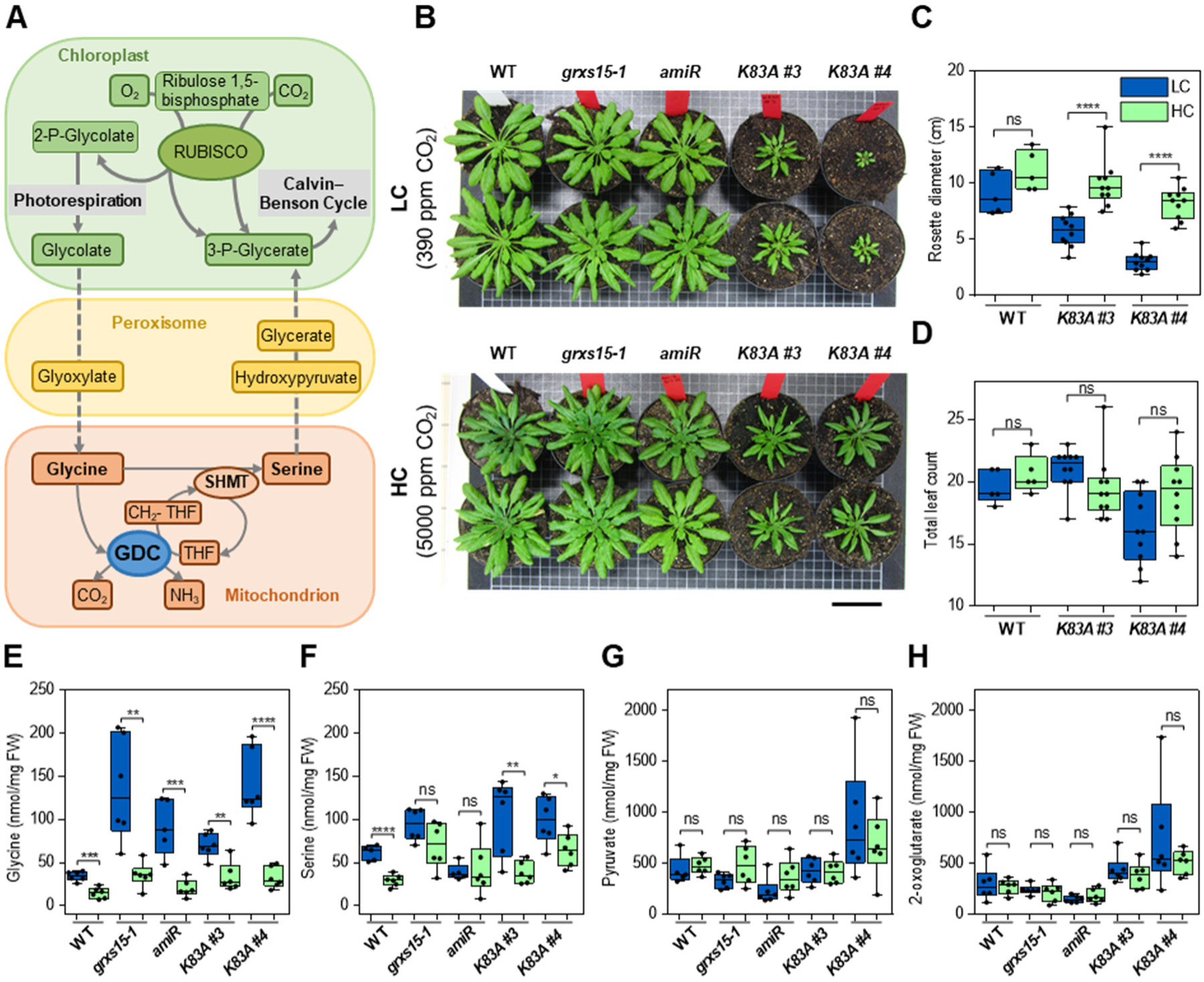
Suppression of the photorespiratory flux enables partial recovery of dwarf *grxs15* mutants. A, Schematic representation of the photorespiratory pathway highlighting the key role of glycine decarboxylase (GDC). For simplicity, the scheme does not indicate the correct stoichiometry. B, Phenotypes of two representative 8-week-old wild type (WT) and *grxs15* mutant plants grown in low CO_2_ (LC, shown in blue) and high CO_2_ (HC, shown in green), respectively, under a 12h/12h day/night regime. Scale bar: 5 cm. C, D, Rosette diameter and total leaf counts of WT and the *K83A* mutants grown in LC or HC (*n* = 5-10). E-H, Effect of atmospheric CO_2_ on the abundance of glycine, serine, pyruvate and 2-oxoglutarate in leaves of 8-week-old WT and *grxs15* mutant plants (*n* = 5-6). All box plots show the median as centre line with the box for the first to the third quartile and whiskers indicating min and max values of the whole data set. Asterisks represent significant differences (*=*P* ≤ 0.1, **= *P* ≤ 0.01, ***= *P* ≤ 0.001, ****= *P* ≤ 0.0001, ns: not significant) calculated according to Student’s *t*-test (α = 0.05). *P*-values: Supplemental Data Set 1.

Growth in 5,000 ppm CO_2_ partially rescued the dwarf phenotype of the most severe *grxs15* mutants *K83A #3* and *#4* compared to plants grown in ambient CO_2_ of 390 ppm (Figure 1, B-D). Diminished RubisCO-mediated oxygenation of ribulose-1,5-bisphophate in plants exposed to high CO_2_ consistently led to a decrease in glycine levels. While this effect is already evident in wild type (WT) plants, it is far more pronounced in *grxs15* mutants, which all accumulate glycine 2 to 4-fold with respect to the WT under ambient conditions. High CO_2_ levels caused glycine to drop to levels comparable to the glycine content in WT plants (Figure 1E). Although serine was also slightly increased in most mutant lines (Figure 1F), the increase in glycine was more pronounced and, thus, consistently led to an increase in the glycine-to-serine ratio, which serves as a measure of impaired GDC activity. Interestingly, a significant decrease in serine was observed in *K83A* mutants grown in high CO_2_ (Figure 1F). This may suggest that especially in rosette leaves other pathways in different compartments contribute to serine biosynthesis (Ros et al., 2014; Anoman et al., 2019). *K83A* mutants also showed increased levels of the photorespiratory intermediates glyoxylate and glycerate in normal air, which was more pronounced in the most severe *K83A #4* mutant. At least in the *K83A* mutants high CO_2_ caused a decrease of these metabolites (Supplemental Figure S2). The respective changes may at least in part explain the severe growth phenotypes. Pyruvate and 2-OG that had been found increased in young *K83A* seedlings (Moseler et al., 2021) were barely altered in rosette leaves and not significantly affected by high CO_2_ (Figure 1, G and H).

### Deletion of the highly abundant ACO3 suppresses the dwarf phenotype of mutants with diminished GRXS15 capacity

Lipoyl synthase, which catalyses the last step in the lipoylation pathway, is a low-abundant mitochondrial protein. Based on the low copy number of only 85 LIP1 proteins per mitochondrion versus more than 10,000 for the [4Fe–4S] cluster proteins aconitase 2 and 3 (ACO2 and ACO3) (Fuchs et al., 2020), we reasoned that LIP1 may not be supplied with sufficient [4Fe–4S] clusters if a bottleneck occurs in the upstream supply chain. In *grxs15* mutants, expression of *ACO2* and *ACO3* were increased up to 2.7-fold and 4-fold, respectively, while expression levels of *LIP1* did not change (Supplemental Figure S3). Even if changes in expression levels would only partially correlate with changes in protein abundance, it is likely that this transcriptional activation further shifts the ACO3/LIP1 abundance imbalance towards ACO3 in *grxs15* mutants. More ACO3 and the respective need for [4Fe–4S] would most likely render LIP1 even less competitive for receiving sufficient clusters to attain catalytic activity. This situation likely aggravates if the auxiliary [4Fe–4S] cluster gets sacrificed for supply of two sulfhydryl groups to an octanoyl residue in each catalytic cycle similar to the mechanism of the bacterial homologue of LIP1, LipA (McCarthy and Booker, 2017). Therefore, we hypothesized that alterations in demand for [4Fe–4S] clusters through modulation of apoprotein abundance may allow to redirect the flux of [4Fe–4S] clusters towards LIP1. Because *aco3* null mutants show only a mild growth impairment compared to WT plants (Figure 2, A and B), and because ACO3 is more abundant than ACO2 (Moeder et al., 2007; Fuchs et al., 2020), we selected the *aco3* mutant for crossing with different *grxs15* mutants (Supplemental Figure S4).

**Figure 2.**
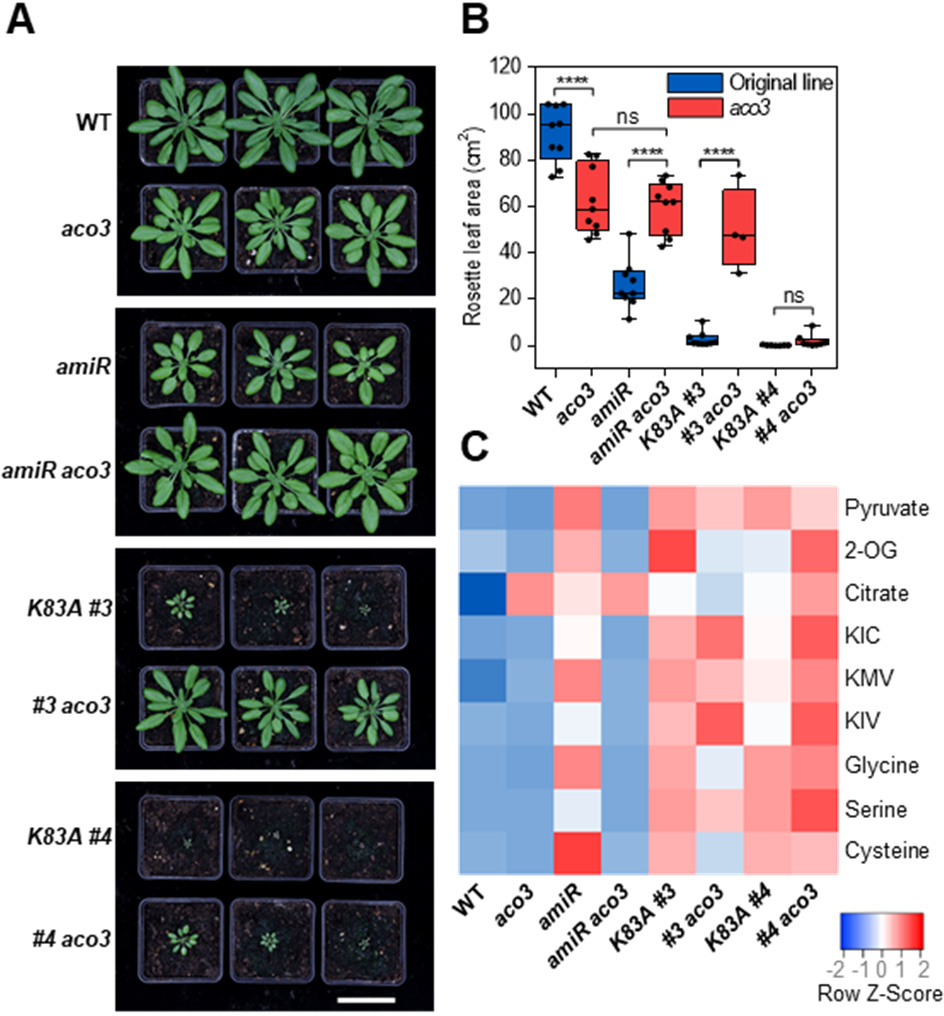
Loss of *ACO3* partially suppresses the dwarfism of *grxs15* mutants. A, 5-week-old plants grown on soil under long-day conditions. *aco3* and double homozygous crosses are compared with WT and the respective parental lines. Scale bar: 5 cm. B, Analysis of rosette leaf area of 4-week-old plants grown on soil (*n* = 4-9). The box plot shows the median as centre line with the box for the first to the third quartile and whiskers indicating min and max values of the whole data set. Asterisks represent significant differences (****= *P* ≤ 0.0001, ns: not significant) calculated according to one-way ANOVA with Tukey’s multiple comparisons test (α = 0.05). *P*-values: Supplemental Data Set 2. Data for WT and the different *grxs15* mutants are shown in blue, data for *aco3* and the respective crosses are shown in red. C, Heat map of key metabolites known to be associated with lipoyl cofactor-dependent enzymes. Metabolites were measured in 8-day-old seedlings (see Supplemental Figure S6 and Supplemental Data Set 2 for absolute values). Z-scores of the mean were calculated for each metabolite and are presented as a heat map. Decreased metabolites are depicted in blue and increased metabolites in red.

Interestingly, loss of *ACO3* in all cases partially suppressed the dwarf phenotypes of *amiR*, *K83A #3* and *#4*, especially at later developmental stages (Figure 2, A and B and Supplemental Figure S5). In both, *amiR* and *K83A #3,* loss of *ACO3* ultimately led to phenotypes similar to *aco3*. For the *amiR aco3* double mutant the root length was already 7.3-fold longer (*amiR aco3* 3.9 ± 0.07 cm versus *amiR* 0.53 ± 0.02 cm; Supplemental Figure 5, A and B) and the rosette of 4-week-old plants were twice as large as in the original *amiR* single mutant (*amiR aco3* 60.0 ± 3.8 cm^2^ versus *amiR* 26.3 ± 3.5 cm^2^; Figure 2, A and B). Ten days after germination, the cross of *K83A #3* and *aco3* (for simplicity hereafter called *#3 aco3*) showed only a tendency for increased root length compared to the *K83A #3* background, but after 17 days, the suppression of the short root phenotype was apparent (Supplemental Figure 5, A-C). When grown on soil for four weeks *#3 aco3* double mutant plants had an 18-fold larger rosette area than the original *K83A #3* mutant (*#3 aco3* 49.9 ± 8.8 cm^2^ versus *K83A #3* 2.7 ± 1.1 cm^2^; Figure 2, B and C). The growth-promoting effect of *ACO3* loss was also apparent in *#4 aco3*, although less pronounced than in *#3*, most likely because *#4* plants are generally more compromised than *#3* plants. While flower stalks of *K83A #3* and *#4* remained short compared to WT, inflorescence development in 8-week-old plants was at least partially rescued by the loss of *ACO3* (Supplemental Figure S5D).

To further characterize the metabolic consequences of the combined decrease in GRXS15-based supply of [2Fe–2S] clusters and loss of *ACO3*, we analysed key metabolites that were all found to be increased in *grxs15* seedlings due to diminished LIP1 activity (Moseler et al., 2021). Notably, accumulation of pyruvate, 2-OG, α-keto acids, and glycine in *amiR* was reverted to *aco3* levels in *amiR aco3* (Figure 2C and Supplemental Figure 6, A-G). Contrary to the expectation that GDC deficiency in *amiR* should cause a decrease in serine as a product of glycine decarboxylation, serine also increased when GRXS15 activity was diminished. Yet, the increased glycine-to-serine ratio consistently points at GDC deficiency, which normalized when *ACO3* was lost. This picture was less consistent in line *K83A #3 aco3*, which showed a decrease in glycine and 2-OG, but no change in pyruvate and the α-keto acids α-ketoisocaproic acid (KIC) and α-keto-β-methylvaleric acid (KMV). α-ketoisovaleric acid (KIV) was even further increased in *K83A #3 aco3*. *K83A #4 aco3* mutants showed an increase in all α-keto acids and 2-OG, but no pronounced change in pyruvate, glycine and serine. Compared to the WT, citrate was increased in *aco3* and all *grxs15* mutants but without any additive effect in double mutants (Figure 2S and Supplemental Figure 6C). With multiple metabolic connections between different metabolites within one compartment or even across different compartments, it can be expected that a metabolic block in one specific enzyme may have rather pleiotropic consequences in several other metabolite pools. This is highlighted by an increase in cysteine in *grxs15* mutants, which trails the increase in the precursors glycine and serine, albeit at much lower concentrations (Figure 2C, Supplemental Figure S6, G-I). Loss of *ACO3* in *amiR* and *K83A #3* caused a decrease in cysteine. Only in the most severe mutant *K83A #4,* additional loss of *ACO3* did not bring down the levels of these three amino acids although the severe growth phenotype was already slightly suppressed (Figure 2, Supplemental Figure S6, G-I). Taken together, these results suggest that loss of the highly abundant [4Fe–4S] sink ACO3 enables redirection of [4Fe–4S] clusters to LIP1 to regain at least partial lipoylation activity in *grxs15* mutants indicated by readjustment of metabolite levels.

### Overexpression of *LIP1* rescues severe *grxs15 K83A* mutants but is deleterious for the wild type and *grxs15* knockdown mutants

Having established experimental evidence for putative redirection of [4Fe–4S] clusters from ACO3 to LIP1 in *aco3* mutants, we next hypothesised that also overexpression of *LIP1* may render the respective sink more competitive over other apoproteins for a limited [4Fe–4S] pool. To test this hypothesis, we constructed overexpression vectors for *LIP1* driven by the CaMV 35S promoter (Figure 3A). To validate correct mitochondrial targeting, we added a C-terminal GFP (Figure 3, A and B; Supplemental Figure S7).

**Figure 3.**
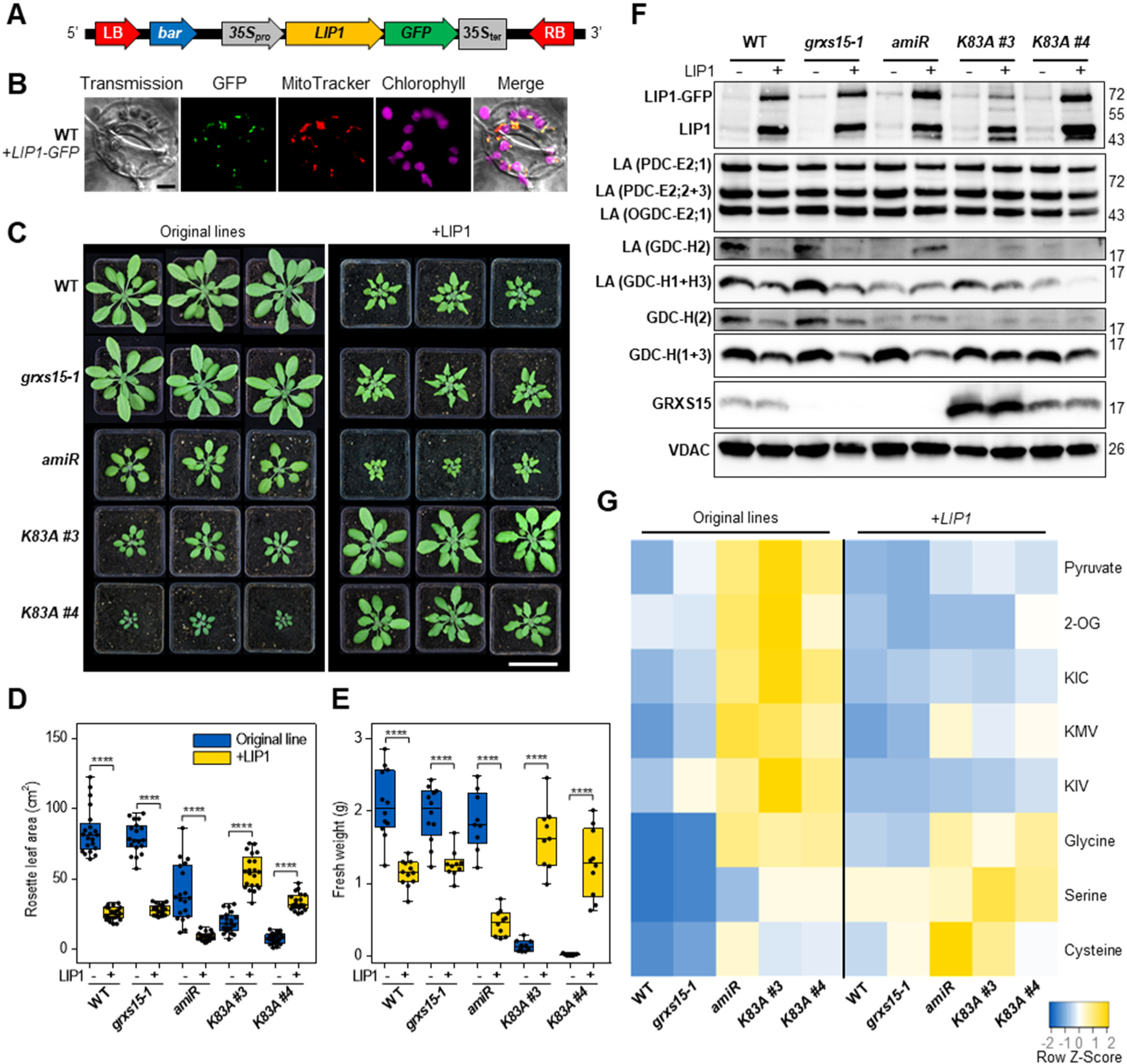
Overexpression of *LIP1* is deleterious for wild type plants and *grxs15* knock-down mutants but beneficial for the *K83A* mutants. A, Schematic representation of the construct used for overexpression of *LIP1*. The expressed protein has GFP fused at its C-terminus. B, Confocal microscopy images of guard cells from 7-day-old wild type seedlings stably expressing *LIP1-GFP*. Fluorescence images were collected with the following wavelength settings: GFP (λ_ex_ = 488 nm; λ_em_ = 520 nm), MitoTracker Orange (λ_ex_ = 543 nm; λ_em_ = 597 nm), chlorophyll autofluorescence (λ_ex_ = 633 nm; λ_em_ = 675 nm). The merge image shows the overlap of all four individual channels. Scale bar = 5 µm. C, 5-week-old plants grown on soil in long-day conditions. Subpanels on the left show the original lines, with their respective genetic background. Subpanels on the right show the same lines overexpressing *LIP1-GFP*. Scale bar: 5 cm. D, Rosette leaf area analysis of 4-week-old plants (*n* = 21). E, Shoot fresh weight of 7-week-old plants (*n* = 9-12). All box plots show the median as centre line with the box for the first to the third quartile and whiskers indicating min and max values of the whole data set. Asterisks represent significant differences (****= *P* ≤ 0.0001) calculated according to one-way ANOVA with Tukey’s multiple comparisons test (α = 0.05). *P*-values: Supplemental Data Set 3. Data for WT and the different *grxs15* mutants are shown in blue, data for the respective lines overexpressing *LIP1* are shown in yellow. F, Protein gel blot analysis with primary antibodies raised against LIP1, GRXS15, LA, GDC-H, and VDAC for loading control. Mitochondria were isolated from seedlings grown in hydroponic culture or from leaves of plants grown on soil and 15-20 µg protein loaded per lane. A blot with extended exposure for the detection of GRXS15 indicates some residual amounts of GRXS15 in *grxs15-1* and *amiR* (Supplemental Figure S1). G, Heat map of metabolites known to be substrates of LA-dependent enzymes or in case of serine and cysteine are derivatives of glycine. Metabolites were extracted from 8-day-old seedlings (for absolute values see Supplemental Figure S15 and Supplemental Data Set 3). Z-scores of the mean were calculated for each metabolite and are presented as a heat map, with decreased metabolites coded in blue and increased metabolites depicted in yellow.

Consistent with our hypothesis, overexpression of *LIP1* largely suppressed dwarfism of *K83A #3* and *#4* in several independent lines. This effect was apparent at all developmental stages with significant increases in root length, rosette size, biomass and flower stalk height (Figure 3, C-E; Supplemental Figures S8-S10). In all independent lines, plants with the *K83A #4* background remained slightly smaller than *K83A #3*. Unexpectedly, however, *LIP1* overexpression had the opposite effect in the WT and the two knockdown mutants *grxs15-1* and *amiR*, where increased LIP1 abundance resulted in significantly reduced growth and delayed development. In these cases, the rosette area of 4-week-old plants was decreased by 70 % for the WT, 64 % in *grxs15-1* and 77 % in *amiR*, which was largely mirrored in the fresh weight of the respective rosettes (Figure 3, C-E). The rosette leaves in these plants showed a curly phenotype similar to other mutants with disturbed mitochondrial metabolism (Supplemental Figure S11; (Lee et al., 2021)). Flower initiation in these mutants was delayed by one to two weeks and the inflorescences remained much shorter than in plants without overexpression of *LIP1* (Supplemental Figure S10C). Notably, parallel experiments in which *LIP1* was overexpressed without *GFP* led to similar results indicating that the GFP tag did not provoke any deleterious effect (Supplemental Figure S12).

To further substantiate either beneficial or deleterious effects of *LIP1* overexpression in different genetic backgrounds, we tested for LIP1 protein abundance with immunodetection in protein blots. Overexpression of *LIP1-GFP* in all lines caused high abundance of LIP1-GFP with the expected molecular mass of ∼70 kDa (Figure 3F, Supplemental Figure S13A). In addition to the LIP1-GFP band, blots probed with α-LIP1 showed another band with an apparent molecular mass of ∼45 kDa likely representing free LIP1, and a distinct band for free GFP when probed with α-GFP (Supplemental Figure S14). The absence of residual fluorescence in the cytosol (Supplemental Figure S7) strongly suggests that the cleavage of LIP1-GFP occurred during or after protein extraction. Although overexpression generally resulted in high abundance of LIP1, some differences in protein levels were apparent. Notably, the LIP1-GFP band at ∼70 kDa was most intense for *amiR* and *K83A #4* but far less intense for *K83A #3*. The lipoylation pattern detected by probing the protein blots with an LA-specific antibody did not change for E2 subunits of PDC and OGDC. Contrary to our expectation, for H1 and H3 subunits of GDC, a slight decrease in lipoylation was found in WT and *grxs15-1* after *LIP1* expression. For H2 in these lines, the decrease was even more pronounced. However, an increased protein lipoylation was observed for the H2 subunit of GDC at least in *amiR* and *K83A #3* overexpressing *LIP1-GFP* (Figure 3F; Supplemental Figure S13). In mitochondria isolated from WT and *grxs15-1* overexpressing *LIP1-GFP*, this was not observed, but it should be noted that the amount of H2 was already diminished in these severely compromised dwarf mutants.

Overexpression of *LIP1* affected the metabolite signature with pronounced decreases of the key metabolites pyruvate, 2-OG and all three BCKAs that generally accumulated in *grxs15* seedlings (Figure 3G and Supplemental Figure S15). For glycine and serine, the picture was less uniform with glycine being increased in WT and *grxs15-1* seedlings overexpressing *LIP1*, but only minor or no changes in the more severe *grxs15* mutants, which all had increased glycine levels without additional LIP1 (Figure 3G and Supplemental Figure S15G). Serine was increased after *LIP1* overexpression in WT and all *grxs15* mutants. Interestingly, all lines except *K83A #4* overexpressing *LIP1* showed an increase in cysteine of 60-100 % compared to control plants (Supplemental Figure S15H). Accumulation of glycine and serine was also found in rosette leaves of soil-grown plants with diminished GRXS15 activity (Figure 1, E and F; Supplemental Figure S16). In these older plants, *LIP1* overexpression caused an increase in glycine and serine, which was largely abolished in high CO_2_. This result suggests that photorespiration was the main source of the increase in glycine, serine and potentially cysteine rather than the overexpression of *LIP1* itself. Different from the positive effect of high CO_2_ on the *K83A* mutants (Figure 1B), the same mutants rescued by *LIP1* overexpression did not show further growth improvement as seen before but rather a decrease in biomass (Supplemental Figure S16A).

### LIP1 causes a dose-dependent phenotypic response and induction of AOX

Next, we wondered why in addition to reduced GDC-H lipoylation overexpression of *LIP1* is deleterious for the WT and for plants with diminished abundance of wild type GRXS15, i.e. *grxs15-1* and *amiR*. While searching for plants with homozygous *LIP1* overexpression in the WT background it became apparent that severity of the phenotype depends on the zygosity with homozygous plants being most severely affected (Figure 4, A-C; Supplemental Figure S17). The zygosity-dependent negative gene dosage effect was also seen in the other *grxs15* mutants (Supplemental Figure S18).

**Figure 4.**
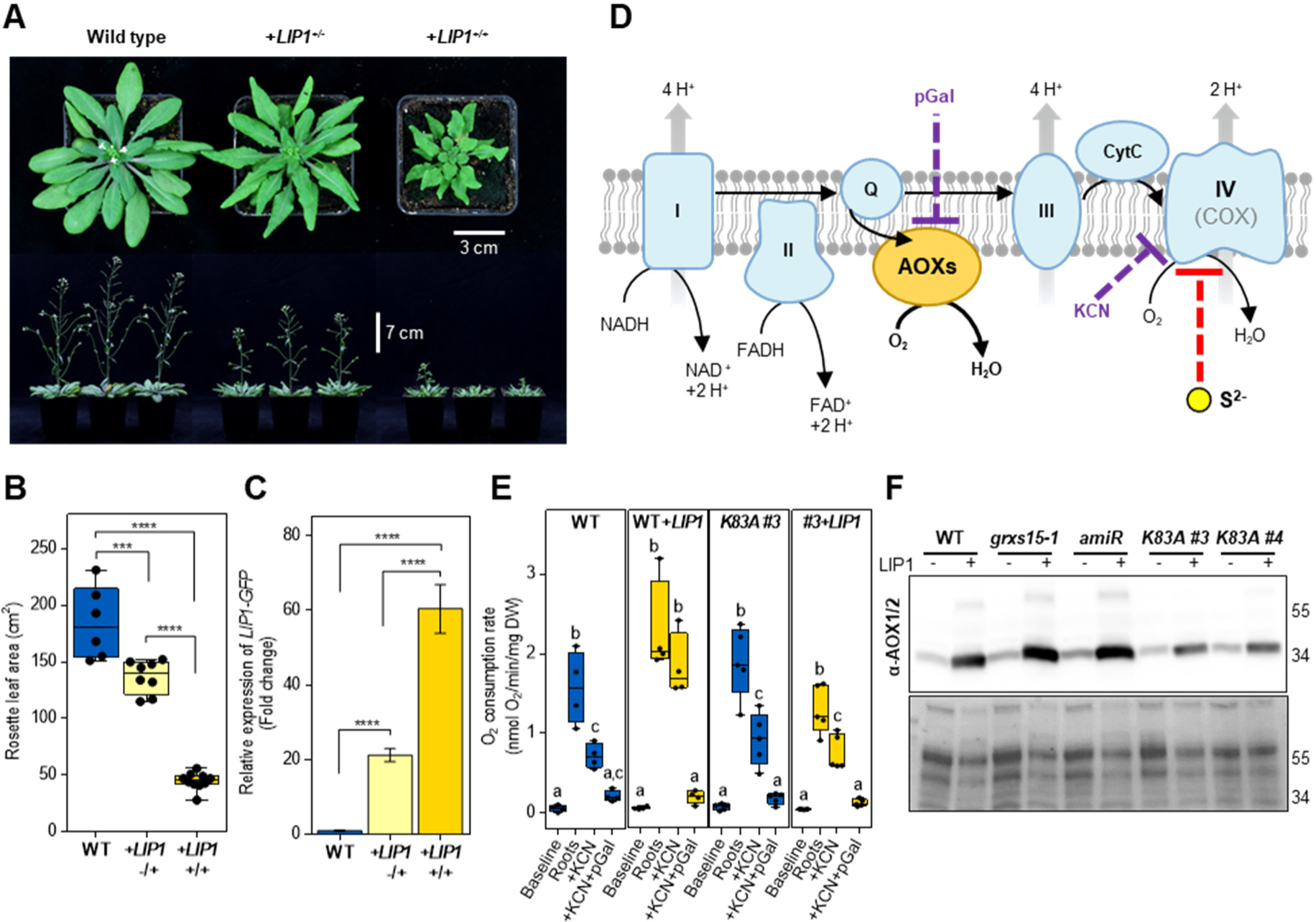
Excessive LIP1 activity causes inhibition of cytochrome c oxidase. A, Representative picture of 6-(top row) and 7-week-old (bottom row) wild type plants overexpressing *LIP1-GFP* in hemizygous (+/−) and homozygous (+/+) form. B, Analysis of rosette area of 4-week-old plants grown on soil in long-day conditions (*n* = 6-8). C, Relative expression level of *LIP1* determined by qRT-PCR. RNA was isolated from 8-day-old seedlings. The presented data are means ± SEM (*n* = 3). The box plot shows the median as centre line with the box for the first to the third quartile and whiskers indicating min and max values of the whole data set. Asterisks represent significant differences (***= *P* ≤ 0.001, ****= *P* ≤ 0.0001) according to one-way ANOVA with Tukey’s multiple comparisons test (α = 0.05). *P*-values: Supplemental Data Set 4. Data for WT and the different *grxs15* mutants are shown in blue, data for the respective lines overexpressing *LIP1* are shown in yellow. D, Scheme of the mitochondrial electron transport chain where alternative oxidases (AOXs) (in yellow) act as terminal electron acceptors when the cytochrome c oxidase (COX) is inhibited by sulfide (S^2−^) (dashed red line). Inhibition of COX by potassium cyanide (KCN) and AOX by propyl gallate (pGal) is indicated by dashed purple lines. E, Respiration of roots of 16-day-old seedlings grown on ½ MS vertical agar plates. After recording the basal root respiration without inhibitors (roots only), COX was first inhibited by addition of KCN at a final concentration of 4 mM. Subsequently, AOXs were inhibited by addition of pGal at a final concentration of 200 µM (*n* = 4-5). Different letters indicate significant differences between treatments calculated according to one-way ANOVA with Tukey’s multiple comparisons test separately for each line (α = 0.05). *P*-values: Supplemental Data Set 4. F, Protein gel blot analysis with primary antibodies raised against AOXs (detects all isoforms: AOX1a-AOX1d and AOX2). Mitochondria were isolated from 16-day-old seedlings grown in hydroponic culture and 15 µg protein loaded per lane. The Amido Black stained membrane at the bottom serves as loading control.

Based on *in vitro* studies with bacterial LipA, it has been suggested that lipoylation activity sacrifices the auxiliary [4Fe–4S] cluster to provide sulfur atoms required for lipoic acid biosynthesis (McCarthy and Booker, 2017). We therefore hypothesised that increased LIP1 activity leads to consumption of more [4Fe–4S] clusters for abstraction of sulfur atoms and that the remains of these clusters fall apart with the inevitable release of sulfide. Inhibition of COX by sulfide would then lead to increased expression and activity of alternative oxidase (AOX) as an overflow valve for electrons in the mETC (Figure 4D; (Selinski et al., 2018)). Consistent with this hypothesis, the COX inhibitor potassium cyanide (KCN) largely blocked respiration in wild type roots but far less in roots from WT overexpressing *LIP1* (Figure 4E). Conversely, *LIP1* overexpression in WT plants resulted in KCN-insensitivity but more pronounced inhibition of respiration when propyl gallate (pGal) as an inhibitor of AOX was applied (Figure 4E; Supplemental Figure S19). The basic respiration of WT plants overexpressing *LIP1* was consistently higher than the respiration of non-transformed plants (Supplemental Figure S19C). Interestingly, respiration of roots from *K83A #3* plants overexpressing *LIP1* decreased significantly in the presence of KCN. With this, the sensitivity of the mETC in *K83A #3* with LIP1 to respiratory inhibitors was more similar to the WT than to WT plus LIP1, which suggests that the *K83A* mutation generates a major bottleneck for Fe–S cluster supply (Figure 4E; Supplemental Figure S19).

Changes in electron flow after *LIP1* overexpression caused a pronounced increase in AOX protein levels in WT plants and the two knockdown mutants *grxs15-1* and *amiR* overexpressing *LIP1* (Figure 4F; Supplemental Figure S20). In the two *K83A* mutants that were largely (*#3*) or partially (*#4*) rescued by LIP1, AOX was still increased but far less abundant than in the two knockdown lines. The relative abundance of AOX in mature leaves of the two *K83A* mutants with higher levels in *#4* plants matches with the degree of growth retardation, which is also more severe in *#4* than in *#3* (Supplemental Figure S20A; Figure 3C).

### Mitochondrial cysteine biosynthesis in Arabidopsis keeps sulfide released by LIP1 under control

To prevent poisoning of COX by sulfide, evolution exploited two mechanisms for conversion of sulfide to largely innocuous products. Animal cells and most fungi oxidise sulfide by the inner mitochondrial membrane protein SQR, while plants can use sulfide in the mitochondrial matrix for cysteine production by OAS-TL C (Theissen et al., 2003; Birke et al., 2012; Vitvitsky et al., 2021). To assess whether the increased mitochondrial sulfide load in *LIP1* overexpression lines can be countered by increasing the endogenous mitochondrial scavenging capacity for sulfide, we additionally overexpressed *OAS-TL* C (Figure 5A). Consistent with the hypothesis that overexpressed *LIP1* releases toxic amounts of sulfide that exceed the endogenous detoxification capacity, we observed at least partial recovery of the dwarf phenotype observed in *LIP1* overexpressors with more than a 2-fold increment in rosette area (Figure 5, B and C). Moreover, the respiration of WT plants overexpressing both *LIP1* and *OAS-TL C* showed again increased sensitivity to KCN, indicating that the inhibition of COX by sulfide originating from *LIP1* was partially rescued by overexpression of *OAS-TL C* (Figure 5D). Overexpression of *OAS-TL C* itself did not exhibit any harmful effect on the WT (Figure 5, B and D).

**Figure 5.**
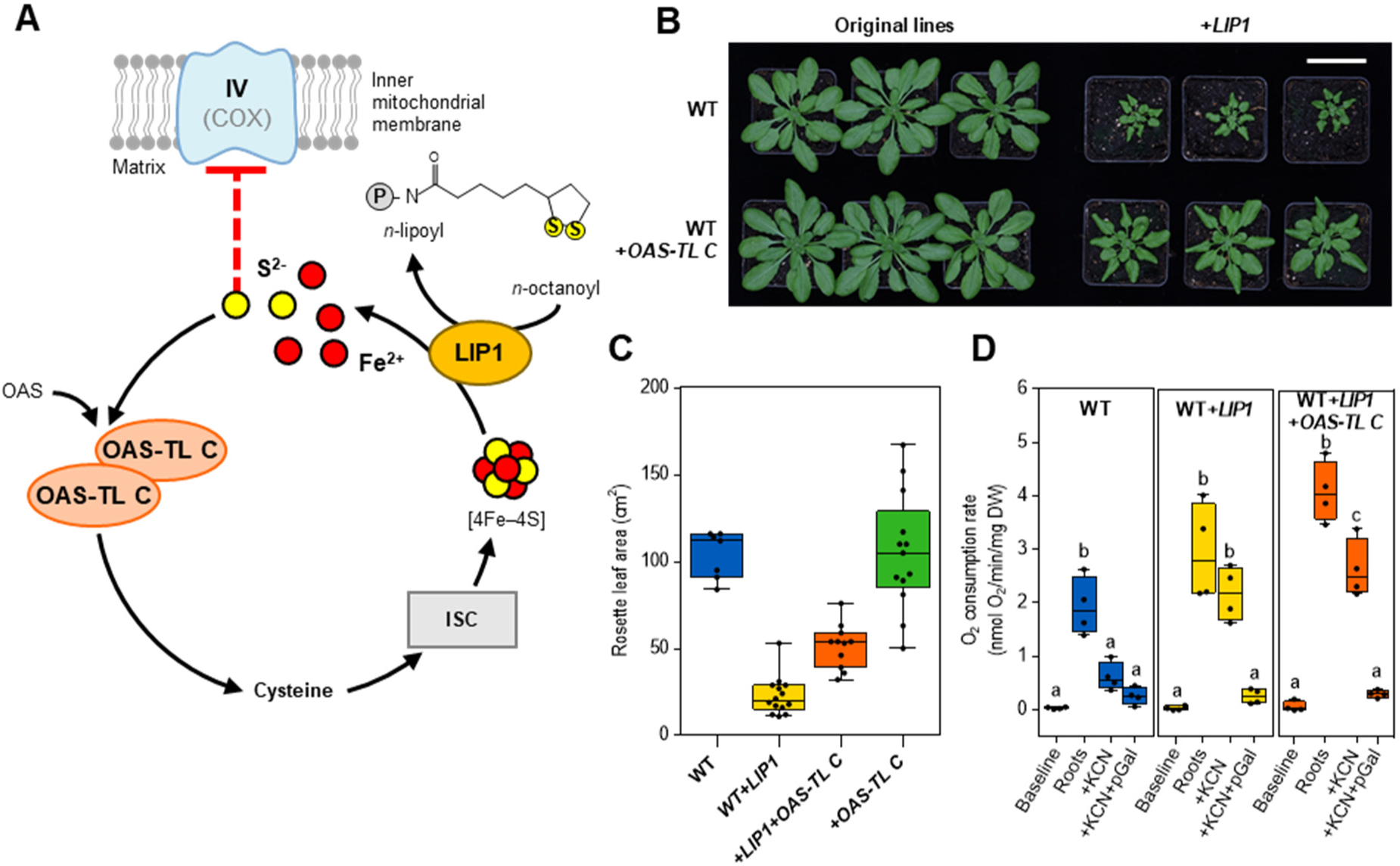
Overexpression of *OAS-TL C* suppresses sulfide toxicity resulting from *LIP1* overexpression. A, Hypothetic model for the lipoylation reaction with concomitant sulfide release and its subsequent refixation by OAS-TL C. Cysteine provides the mitochondrial iron–sulfur cluster (ISC) assembly machinery with the required sulfur. LIP1 sacrifices a [4Fe–4S] cluster and releases sulfide that needs to be detoxified to avoid poisoning of COX. B, Representative photos of 5-week-old plants grown on soil in long-day conditions. Plants overexpressing *LIP1-GFP* (right) are compared with their respective genetic background (left). Scale bar: 5 cm. C, Rosette area analysis of 4-week-old plants overexpressing *LIP1* or *LIP1* plus *OAS-TL C* compared with the WT (*n* = 7-14). D, Respiration of roots of 16-day-old seedlings grown on ½ MS vertical agar plates. After recording the basal root respiration without inhibitors (‘Roots’), COX was first inhibited by addition of KCN at a final concentration of 4 mM. Subsequently, AOXs were inhibited by addition of pGal at a final concentration of 200 µM (*n* = 4). All box plots show the median as centre line with the box for the first to the third quartile and whiskers indicating min and max values of the whole data set. Different letters indicate significant differences between treatments calculated according to one-way ANOVA with Tukey’s multiple comparisons test separately for each line (α = 0.05). *P*-values: Supplemental Data Set 5.

## Discussion

Lipoylation is a rare, but yet essential posttranslational lysine modification that is evolutionarily conserved from bacteria to humans (Rowland et al., 2018). Defects in lipoylation reactions thus cause severe metabolic disorders or even lethality in plants, yeast and mammals (Sulo and Martin, 1993; Yi and Maeda, 2005; Navarro-Sastre et al., 2011; Tort et al., 2013; Mayr et al., 2014; Bauwe, 2023). In plants, deficiencies in Fe–S cluster supply become most apparent from defects in GDC and the concomitant increase in glycine and the glycine-to-serine ratio (Przybyla-Toscano et al., 2022). This increase in glycine can be prevented if the metabolic demand on GDC is minimized by suppressing photorespiration in a high CO_2_ atmosphere (Przybyla-Toscano et al., 2022). The same physiological response occurs also in mutants with severe deficiencies in GRXS15-dependent [2Fe–2S] cluster transfer between the two Fe–S cluster assembly machineries (Figure 1; Figure 6). Without any external input, the lipoylation-deficient phenotype can also be suppressed by altering the distribution of [4Fe–4S] clusters between the highly abundant ACO3 and the very low abundant LIP1 (Figure 2; Figure 3C). Under normal growth conditions, loss of *ACO3* does not severely compromise the plant because mitochondria also contain ACO2 to serve in the TCA cycle (Hooks et al., 2014). Losing ACO3 as an [4Fe–4S] apoprotein makes more clusters available for LIP1 and thus enables increased LIP1 activity even in severe *grxs15* mutants. As an orthogonal approach, also *LIP1* overexpression rescued the most severely compromised *grxs15* mutants carrying a K83A variant of GRXS15 (Figure 3C). All these results consistently show that LIP1 requires sufficient [4Fe–4S] cluster supply to attain full lipoylation activity. Previously, it was considered that GRXS15 might also be involved in repair of Fe–S clusters or have an oxidoreductase function (Couturier et al., 2015; Zhang et al., 2018). Our data, however, show that at least in severely compromised *grxs15* mutants, cluster supply rather than any other putative function of GRXS15 is the limiting factor for plant performance.

**Figure 6.**
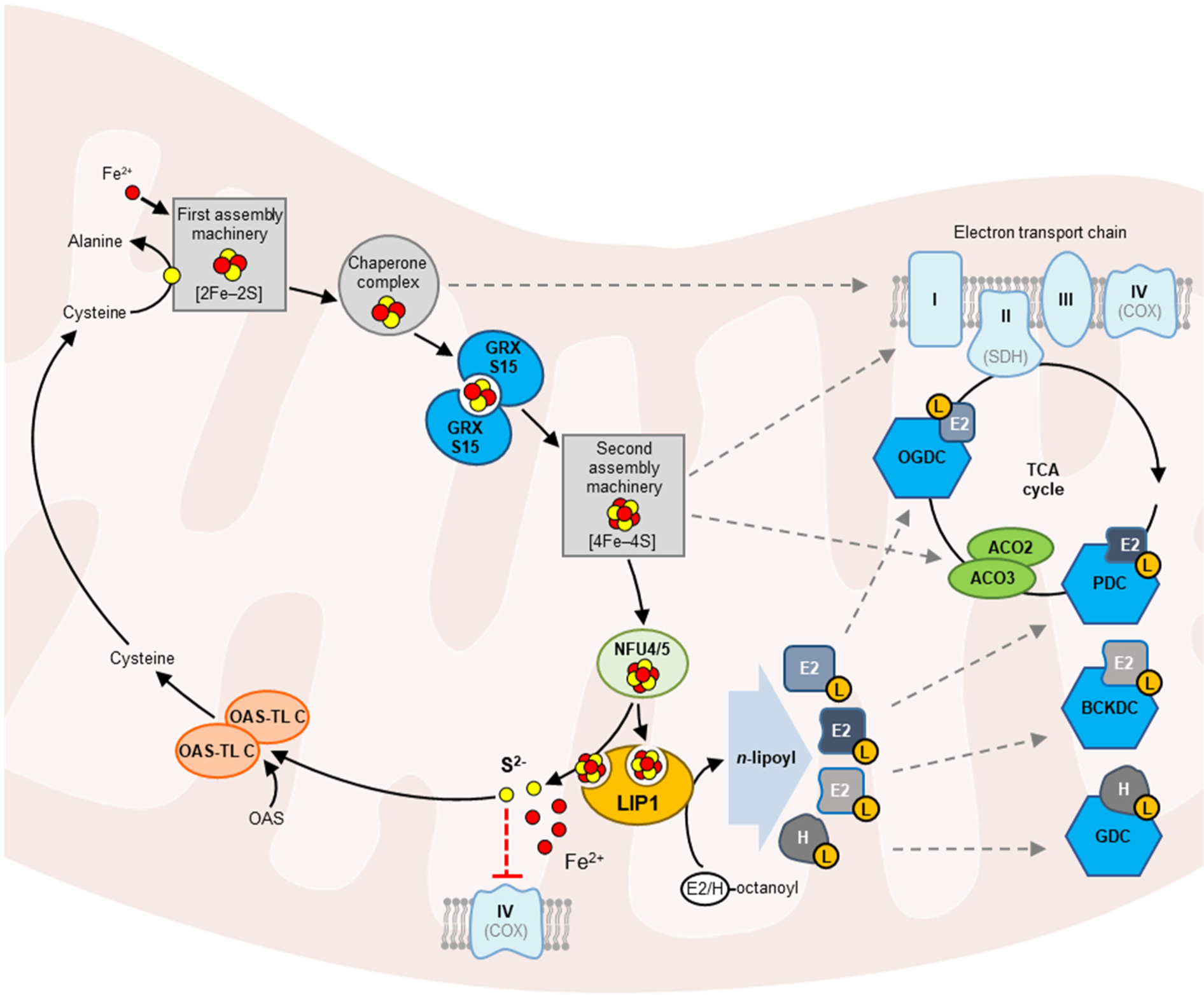
Working model of iron–sulfur cluster assembly and transfer machineries in plant mitochondria, with LIP1 as an Fe–S cluster consuming enzyme. On the top left, a simplification of the first assembly machinery, which utilises cysteine as sulfur source (yellow circles) and iron (red circles) to build [2Fe– 2S] clusters, is depicted. After assembly, a chaperone complex acts as a hub for further direct or indirect delivery of [2Fe–2S] clusters to target proteins such subunits of complexes I, II and III in the mitochondrial electron transport chain (ETC) (dashed arrow) and to glutaredoxin S15 (GRXS15). GRXS15 is the key enzyme for transfer of [2Fe–2S] clusters from the first assembly machinery to the second, where [4Fe–4S] clusters are being built. These are then distributed to target proteins, such as ETC complexes, aconitases 2 and 3 (ACO2/3) and lipoyl synthase (LIP1). Delivery of [4Fe–4S] to LIP1 relies on the NifU-like proteins NFU4/5. LIP1 requires one [4Fe–4S] cluster for its catalytic activity and uses a second, auxiliary, cluster as sulfur source to synthesise the lipoyl cofactor, which is necessary for the activity of four mitochondrial dehydrogenase complexes glycine decarboxylase (GDC), pyruvate dehydrogenase (PDC), 2-oxoglutarate dehydrogenase (OGDC), and branched-chain α-keto acid dehydrogenase (BCKDC). LIP1 transfers two sulfur atoms from the auxiliary cluster to the *n*-octanoyl residue on the dehydrogenase subunits E2 or H to convert these into *n*-lipoyl prosthetic groups (L). In the absence of any established repair mechanism, the remnants of the cluster are assumed to disintegrate and release sulfide. Similarly, entire auxiliary clusters may fall apart if they are not used immediately, *e.g.*, in case of substrate shortage for LIP1 or increased LIP1 abundance. To prevent poisoning of cytochrome *c* oxidase (COX, complex IV), the toxic sulfide is utilised by *O*-acetylserine-(thiol)-lyase C (OAS-TL C) to synthesise cysteine, which can be reused for a new round of Fe–S cluster assembly.

Wild-type plants and mutants less compromised in [2Fe–2S] cluster transfer, however, showed a very different response to *LIP1* overexpression (Figure 3C). The very low abundance of endogenous GRXS15 in *grxs15-1* and *amiR* plants caused a partial photorespiratory phenotype identified by increased glycine-to-serine ratios already. Apparently, however, the capacity for [2Fe–2S] cluster transfer is still sufficient to enable a minimum [4Fe–4S] cluster supply to LIP1 to avoid accumulation of more toxic photorespiratory intermediates upstream of glycine (Timm et al., 2008; Dellero et al., 2016). Overexpression of *LIP1* in these mutants and in the WT allows for efficient redirection of more [4Fe–4S] clusters to LIP1, which became most apparent by the reversion of the photorespiratory metabolite signature in the *grxs15* knockdown mutants. Yet, increased *LIP1* levels in these plants surprisingly turned out to be deleterious. Dependence of this phenotype on LIP1 is also supported by a gene-dosage-dependent increase in severity in hemizygous and homozygous *LIP1* overexpressors. Apparently, increased abundance of LIP1 can attract more [4Fe–4S] clusters than what would normally be necessary to maintain physiological needs, which ultimately causes toxic side effects (Figure 6).

Based on *in vitro* experiments with recombinant LipA from *E. coli* and human LIAS (both homologs of Arabidopsis LIP1), it was reported that the enzymes in the presence of NFUs can potentially direct all four sulfide ions from the auxiliary cluster into the lipoyl product rather than releasing two of them into solution (McCarthy and Booker, 2017; Warui et al., 2022). Yet, if only a single catalytic cycle is possible because NFU is missing, only two sulfur atoms are extracted from a [4Fe–4S] cluster while the residuals fall apart and release the remaining two sulfides. Detection of sulfide is a major challenge as all current techniques have severe disadvantages and limitations (Kolluru et al., 2013). This applies even more when specific measurements of the sulfide pool in the mitochondrial matrix are to be conducted. Here, we thus indirectly deduce that overexpression of *LIP1* goes along with a pronounced increase in free sulfide. First, COX was strongly inhibited in *LIP1* overexpression plants. With an IC_50_ as low as 6.9 nM (Birke et al., 2012) already a minor increase in sulfide would cause inhibition of COX (Figure 4). This inhibition, possibly in combination with reductive stress imposed by sulfide, triggers a retrograde signal leading to induction of AOX (Fuchs et al., 2020). Increased AOX may compensate for loss of COX activity by offering an alternative route for transfer of electrons to molecular oxygen to avoid overreduction of the ubiquinone pool and concomitant production of reactive oxygen species in the mETC (Millar et al., 2011; Selinski et al., 2018). The predominant isoform AOX1a has been shown to be involved in dissipating thiol-mediated reductive stress (Fuchs et al., 2022). Whether the sulfide released by LIP1 accumulates to sufficient levels for generation of reductive stress remains unknown at this stage. The increased respiration observed under these circumstances in detached Arabidopsis roots is likely an attempt of plants to provide sufficient energy in form of ATP even with a truncated mETC. Because this necessarily goes along with an increased carbon turnover, the block of respiration by sulfide has severe bioenergetic consequences and ultimately causes growth retardation (Birke et al., 2012).

The second piece of evidence for release of sulfide by LIP1 in Arabidopsis results from suppression of the retarded phenotype by overexpressing *OAS-TL C* (Figure 5B). To prevent accumulation of sulfide in the matrix, plants contain local cysteine biosynthesis capacity in mitochondria (Haas et al., 2008; Heeg et al., 2008; Álvarez et al., 2012b). Overexpression of *OAS-TL C* suppresses the dwarf phenotype of WT plants overexpressing *LIP1*. Taken together, all these results support the conclusion that [4Fe–4S] clusters inserted in LIP1 collapse and release poisonous sulfide. Based on the finding that in the absence of NFU the auxiliary [4Fe–4S] cluster collapses after extraction of two sulfur atoms (McCarthy and Booker, 2017) one may speculate whether sulfide toxicity is a consequence of such cluster disintegration during catalysis.

Increased enzymatic activity of overexpressed *LIP1* in Arabidopsis would require the respective substrates for lipoylation. Lipoyl synthases use the octanoylated GDC subunit H2 as their primary substrate for synthesis of the lipoyl moiety, which is subsequently transferred to E2 subunits of the other dehydrogenase complexes (Schonauer et al., 2009; Solmonson and DeBerardinis, 2018; Pietikäinen et al., 2021; Bauwe, 2023). The overall lipoylation pattern on the respective proteins, however, was not massively altered in *LIP1* overexpression lines, especially in WT and GRXS15 knockdown lines. The need for a suitable substrate for the overexpressed LIP1 thus leaves a conundrum. One possibility for providing new substrates for lipoylation would be the presence of lipoamidase activities similar to the sirtuin lipoamidase activities found in mammals and *E. coli* (Mathias et al., 2014; Rowland et al., 2017). So far, however, no such activity has been reported for plants.

An alternative explanation for the acute toxic effects resulting from overexpression of lipoyl synthases may result from cluster instability. The [4Fe–4S] cluster of LipA is known to be sensitive to traces of oxygen and for losing iron and sulfide to yield a [2Fe–2S] cluster upon exposure to air (Ollagnier-de Choudens et al., 2000). Especially in situations with limited substrate supply for the lipoylation reaction, it cannot be excluded that lipoyl synthases lose [4Fe–4S] clusters through oxidative decomposition. If the remaining [2Fe–2S] cluster also becomes unstable after its replacement by a new [4Fe–4S] cluster, this process would ultimately set free four sulfides and thus generate a severe sulfide load in the matrix. Oxidative decomposition of clusters from overexpressed *LIP1* suggests that this likely does happen from the endogenous protein as well, particularly when only limited substrate is available. Interestingly, Arabidopsis mutants deficient in mitochondrial acyl carrier proteins have a similar phenotype as *LIP1* overexpressors. While this is at least in parts caused by lipoylation deficiency (Fu et al., 2020), it might be possible that such mutants also suffer from increased sulfide and inhibition of COX.

The pronounced sulfide release after overexpression of LIP1 in WT plants suggests that the mitochondrial Fe–S cluster assembly machineries have sufficient capacity to serve all available [4Fe–4S] apoproteins. The uncontrolled release of sulfide as a potentially toxic side product suggests that tight control of protein abundance might be the only means for risk management. This is consistent with low abundance of LIP1 in Arabidopsis suspension culture cells (Fuchs et al., 2020) and the absence of any evidence for induction of *LIP1* under specific stress situations. Low abundance of lipoyl synthases is also found in *Saccharomyces cerevisiae* (Ho et al., 2018) and in human cells where LIAS is present with only 23,904 copies per cell, which represents just 0.007 % of all mitochondrial proteins (Morgenstern et al., 2021). Uncontrolled potential release of sulfide by lipoyl synthases also provides a strong argument for maintaining sulfide detoxification systems in mitochondria of most eukaryotic lineages (Theissen et al., 2003; Kabil and Banerjee, 2010; Birke et al., 2012).

Irrespective of whether the released sulfide is a side product of the [4Fe–4S] cluster-sacrificing catalytic activity of lipoyl synthases or a result of uncontrollable oxidative decomposition, the experimental system with overexpression of lipoyl synthases associated with distinct and easily observable phenotypes offers an experimental framework to further study mitochondrial Fe–S assembly and cluster distribution. The increasing severity of phenotypes in mutants with compromised GRXS15-dependent [2Fe–2S] cluster transfer in combination with *LIP1* overexpression and the redirection of [4Fe–4S] clusters through sink modulation both highlight the applicability of the assay. Similarly, one can envisage that the tools can be used to further study mitochondrial sulfide production or detoxification.

## Materials and methods

### Plant material and growth conditions

The *Arabidopsis thaliana* ecotype Columbia-0 ([L.] Heynh.) was used as the main experimental organism. The T-DNA insertion line *grxs15-1* (SALK_112767) has been described earlier (Moseler et al., 2015). Seeds of *aco3* (SALK_014661) (Moeder et al., 2007) were obtained from NASC (Nottingham Arabidopsis Stock Centre, https://arabidopsis.info/). The knockdown line *amiR* (*GRXS15^amiR^* (Ströher et al., 2016)) was provided by Janneke Balk, John Innes Centre, University of East Anglia, UK. The complementation lines *K83A #3* and *K83A #*4 (*grxs15-3 UBQ10_pro_*:*GRXS15 K83A*) were described previously (Moseler et al., 2015; Moseler et al., 2021). The lines *amiR*, *K83A #3* and *#4* were crossed with *aco3* and the double homozygous F_3_ generations were used throughout this study.

For phenotypic characterization of seedlings, seeds were surface-sterilized with 70 % (v/v) ethanol for 10 min followed by washing with sterile water and plated on half-strength Murashige and Skoog medium (½ MS, Duchefa Biochemie, Haarlem, NL) with addition of 0.1 % (w/v) sucrose, 0.05 % (w/v) MES (pH 5.8, KOH) and solidified with 0.8 % (w/v) agar. Seeds were stratified at 4 °C in the dark for 2-3 days. Plates were incubated vertically under long-day conditions (16 h light at 22 °C and 8 h dark at 18 °C) at a light intensity of ∼100 µmol photons m^−2^ s^−1^ and 50 % air humidity. Ten days after germination, root growth was documented photographically and root length was measured using Fiji ImageJ (https://imagej.nih.gov/ij/). For documentation of seedling phenotypes, 17-day-old seedlings were transferred from plates to a black mat and photographed.

For hydroponic seedling culture, surface-sterilized seeds were gently sown on a layer of microagar floating in sterile glass pots filled with 50 mL liquid ½ MS medium with 1 % (w/v) sucrose, 0.04 % (w/v) MES (pH 5.8, KOH). Pots were closed with transparent plastic lids, placed at 4 °C in the dark for two days for seed stratification, and subsequently transferred onto rotary shakers and gently agitated in the growth cabinet under long-day conditions for 16 days.

To obtain larger plants, the seeds were placed onto a standard soil mixture (Floradur B-seed, Perlite Perligran 0-6 and quartz sand in a ratio of 10:1:1, respectively) and stratified for at least 2 days in the dark at 4 °C and high humidity. Subsequently, pots were transferred to the light under long-day conditions (16 h light at 21 °C and 8 h dark at 19 °C) with light intensity of 100-120 µmol photons m^−2^ s^−1^ and 50 % air humidity. The plants were grown in individual pots randomly distributed among greenhouse trays and documented photographically after four weeks for rosette area measurements using Leaf Lab, a custom-written MATLAB script) and after eight weeks for documentation of inflorescences. For determination of the fresh weight, whole rosettes of 7-week-old plants were cut at the base and directly weighted with an analytical scale. To track the floral induction, the growth of plants was monitored three times a week for a visible floral bud. Similarly, the height of the main inflorescence was measured three times a week with a ruler, starting from the base on the rosette. To characterize rosette development and leaf phenotypes, 31-day-old rosettes were dissected and the leaves aligned according to their developmental age.

For experiments with altered CO_2_ conditions, seeds were surface-sterilized with chloric acid and sown on a mixture of soil (Einheitserde; Einheitserdewerk Uetersen Werner Tantau GmbH und Co. KG, Uetersen, DE) and vermiculite (4:1). Pots were incubated at 4 °C for at least two days for stratification and transferred to a growth chamber with defined environmental conditions (SANYO) for eight weeks (principal growth stage 5.1) (Boyes et al., 2001). Plants were grown either in high CO_2_ (HC; 5,000 ppm CO_2_) or in low CO_2_ (LC, 390 ppm CO_2_) for control. In both conditions, the photoperiod was 12/12 h day/night cycle with light intensity of ∼100 µmol m^−2^ s^−1^ and ∼70 % relative humidity. Plants were regularly watered and fertilized weekly (0.2 % Wuxal, Aglukon, Düsseldorf, DE).

For genotyping, a complete list of primers is provided in Supplemental Table S1.

To analyse the zygosity of plants expressing the K83A variant of GRXS15, plants were allowed to self-fertilize. Mature siliques were subsequently checked on a stereomicroscope (Leica M165 FC) for aborted seeds, which would be expected if the transgene was still segregating.

### Generation of transgenic plants

For overexpression of *LIP1* the coding sequence was amplified from cDNA generated from WT plants with primers indicated in Supplemental Table S2 adding *att*B sites. The obtained fragment was purified and BP-recombined into the pDONR207 Gateway donor vector (Invitrogen Waltham, USA). Subsequently, the recombined pDONR207 vector was LR-recombined with a modified pSS01 destination vector (Brach et al., 2009) containing 35S promoter and the gene coding *GFP* in frame with the C-terminus of the Gateway cassette. Alternatively, pDONR207_*LIP1* with a stop codon was LR-recombined with pB7WG2 (Karimi et al., 2002), for overexpression of *LIP1* without *GFP*. After recombination, the expression clone was verified by colony PCR after transformation into *E. coli* DH5α. The vector pMDC32_*35S_pro_*:*OAS-TL C* for overexpression of *OAS-TL C* was kindly provided by Cecilia Gotor (Álvarez et al., 2012a).

*Agrobacterium tumefaciens* (AGL-1) were transformed by electroporation with the final constructs and used to transform plants by floral dipping (Clough and Bent, 1998). T_1_ seeds from plants transformed with pSS01_*35S_pro_*:*LIP1-GFP* and pB7WG2_*35S_pro_*:*LIP1* were harvested and selected either with 240 mg/L glufosinate ammonium solution (Basta) or screened for fluorescence on a stereomicroscope (Leica M165 FC) equipped with a GFP filter (excitation: 470 ± 20 nm, emission: 525 ± 25 nm). T_1_ plants transformed with pMDC32_*35S_pro_*:*OAS-TL C* were screened *in vitro* by 20 µg/mL hygromycin (Harrison et al., 2006). The following generations of mutants were screened by herbicide resistance or fluorescence for segregation, growing mutants in agar plates or on soil. Only lines following a Mendelian segregation were kept (Supplemental Tables S3-S5). Three independent lines for each obtained mutant were selected and screened for the homogeneity of the effects induced by the insertions.

### Subcellular localization of proteins

To verify mitochondrial targeting of LIP1, 7-day-old seedlings stably expressing *LIP1-GFP* were vacuum-infiltrated for 30 min with 200 nM MitoTracker Orange (Invitrogen). To test for colocalization seedlings were imaged on a confocal microscope (Zeiss LSM 780, connected to an Axio Observer.Z1 (Carl Zeiss Microscopy, Jena, DE). Imaging was performed with a C-Apochromat x40 (C-Apochromat 40x/1.2 W corr) water-immersion objective. Fluorescence was recorded with λ_ex_ = 488 nm and λ_em_ = 520 nm for GFP, λ_ex_ = 543 nm, λ_em_ = 597 nm for MitoTracker Orange and λ_ex_ = 633 nm, λ_em_ = 675 nm for chlorophyll autofluorescence. Data acquisition was performed using Zen 3.2 (blue edition) Zeiss software.

### Isolation of intact mitochondria

High-pure intact mitochondria were isolated from 16-day-old seedlings grown in liquid culture following the protocol of Escobar et al. (2006) with slight modifications. About 10-20 g seedling material was ground with 2 g quartz sand in 300 mL extraction buffer (250 mM sucrose, 1.5 mM EDTA, 15 mM MOPS, 0.4 % (w/v) BSA, 0.6 % (w/v) PVP-40, 10 mM DTT, 100 mM ascorbic acid) and filtered through a layer of Miracloth (Merck Millipore) for two times. Cell debris was removed by centrifugation at 1,300 *g* for 5 min (Beckman Centrifuge Avanti® J-26-XP). Afterwards, the supernatant was centrifuged at 18,000 *g* for 20 min to obtain a pellet. The pellet was gently resuspended with a fine brush in 300 mL wash buffer (300 mM sucrose, 10 mM TES, 0,1 % (w/v) BSA). Both centrifugation steps were repeated a second time. Resuspended fractions (∼2 mL) were loaded on PVP gradient (300 mM sucrose, 10 mM TES, 0.1 % (w/v) BSA, 32 % (v/v) Percoll, 0-6 % (w/v) PVP-40) and centrifuged at 40,000 *g* for 40 min with disengaged active-deceleration. Mitochondria fraction at the lower part of the gradient was recovered and washed twice in 60 mL final wash buffer (300 mM sucrose, 10 mM TES) reobtaining mitochondria pellet with centrifuged at 23,700 *g* for 15 min and resuspending it by carefully agitating the tube. The mitochondria pellet was finally resuspended in 500 µL final wash buffer.

Leaf mitochondria from older plants were isolated following the “Method A” described in Keech *et al*. (2005) (Keech et al., 2005) for crude, well-coupled mitochondria. About 5 g of 7-week-old leaves from plants grown on soil in long-day conditions were used.

### Protein blotting and immunodetection

After estimation of protein concentrations via Bradford (Roti-Quant) monitoring absorbance at 595 nm with a CLARIOstar microplate reader, 15 µg of pure isolated mitochondria were mixed with Laemmli buffer (2 % (w/v) SDS, 50 mM Tris-HCl pH 6.8, 0.002 % (w/v) bromophenol blue, 5 % (v/v) β-mercaptoethanol and 10 % (v/v) glycerol), sonicated for 2 min in ultrasonic bath (Bandelin Sonorex) and boiled at 95 °C for 5 min. The samples were electrophoresed on sodium dodecyl sulfate–polyacrylamide gel electrophoresis (SDS–PAGE) precast 4-20 % Mini-PROTEAN TGX gel (Bio-Rad) to separate the proteins.

For immunoblotting the separate protein were semi-dry blotted or wet-blotted on BioTrace PVDF Transfer Membrane (Pall Corporation). Blots were probed in TBS-T with 1:1,000 anti-LIP1 (G231072/63, (Moseler et al., 2021)), 1:1,000 anti-LA (ab58724; Abcam), 1:5,000 anti-GDC-H (As05 074; Agrisera), 1:2,500 anti-GRXS15 (Moseler et al., 2015), 1:5,000 anti-GFP (A-6455; Thermo) or 1:1,000 anti-AOX1/2 (AS04 054; Agrisera) antibodies conjugated to horseradish peroxidase (HRP). 1:20,000 Goat anti-rabbit poly-horseradish peroxidase secondary antibody (#31461; Thermo) was used for detection of chemiluminescence and developed using Pierce ECL Plus Western Blotting Solution (Thermo). Signals were detected on INTAS ECL ChemoStar imager. For the loading control, the PVDF membrane is stained with Amido black staining (0.1 % (w/v) amido black, 45% (v/v) ethanol, 10 % (v/v) acetic acid) or a dedicated gel was prepared and proteins were stained with PageBlue Protein Staining solution (Thermo).

### Metabolite analysis

The determination of metabolites related to photorespiration was carried out by liquid chromatography coupled to tandem mass spectrometry (LC-MS/MS) analysis using the LCMS-8050 system (Shimadzu, Japan). For this purpose, we harvested leaf material (∼25 mg) from fully expanded rosette leaves of plants at growth stage 5.1 (Boyes et al., 2001) grown in high CO_2_ (HC - 5000 ppm CO_2_) or low CO_2_ (LC – 390 ppm CO_2_) at the end of the day (EoD, 11 h illumination). The material was immediately frozen in liquid nitrogen and stored at −80 °C until further analysis. Extraction of soluble primary intermediates was carried out using LC-MS grade chemicals according to (Arrivault et al., 2009; Arrivault et al., 2015) and the samples were analysed exactly as described in Reinholdt et al. (2019). The compounds were identified and quantified using multiple reaction monitoring (MRM) according to the values provided in the LC-MS/MS method package and the LabSolutions software package (Shimadzu, JP). Authentic standard substances (Merck, DE) at varying concentrations were used for calibration and peak areas normalized to signals of the internal standard ((morpholino)-ethanesulfonic acid (MES), 1 mg/mL). Data were interpreted using the Lab solution software package (Shimadzu, JP).

Regarding the other metabolites, 40-50 mg of 8-day-old seedlings growing on agar plates in long-day condition were collected at beginning of the day (BoD: 2 h after the light is on) and flash-frozen in liquid nitrogen before powdering with glass beads using a TissueLyser II (Qiagen).

Ion chromatography was performed for absolute quantification of anion content by extraction with 0.7 mL ultra-pure water for 45 min at 95 °C. Samples were centrifuged at 4 °C and 20,000 *g* for 15 min and 100 µL of the supernatant were transferred to a chromatography vial. After addition of 200 µL ultra-pure water (1:3 dilution), anions were separated using an IonPac AS11-HC (2 mm, Thermo Scientific) column connected to an ICS-5000 system (Thermo Scientific) and quantified by conductivity detection after cation suppression (ASRS-300 2 mm, suppressor current 23-78 mA). Prior to separation, the column was heated to 30 °C and equilibrated with five column volumes of solvent A (ultra-pure water) at a flow rate of 0.3 mL min^−1^. Separation of anions was achieved by increasing the concentration of solvent B (100 mM NaOH) in buffer A linearly as follows: 0-8 min: 4 % B, 18-25 min: 4 %-18 % B, 25-43 min: 19 %-30 % B, 43-53 min: 30 %-62 % B, 53-53.1 min: 62 %-80 % B, 53.1-59 min: 80 % B, 59-59.1 min: 4 % B, and 59.1-70 min: 4 % B. Data acquisition and quantification were performed with Chromeleon 7 software (Thermo Scientific).

Free amino acids and thiols were extracted from 40-50 mg of seedlings with 0.4 mL of 0.1 M HCl in an ultrasonic ice-bath for 10 min. The resulting extracts were centrifuged twice for 10 min at 4 °C and 16,400 *g* to remove cell debris. Amino acids were derivatized with AccQ-Tag reagent (Waters) and determined as described in Weger et al. (2016). Total glutathione and cysteine were quantified by reducing disulfides with DTT followed by thiol derivatization with the fluorescent dye monobromobimane (Thiolyte, Calbiochem). Derivatization was performed as described in Wirtz et al. (2004). UPLC-FLR analysis was carried out using an Acquity H-class UPLC system. Separation was achieved with a binary gradient of buffer A (100 mM potassium acetate, pH 5.3) and solvent B (acetonitrile) with the following gradient: 0 min 2.3 % buffer B, 0.99 min 2.3 %, 1 min 70 %, 1.45 min 70 %, and re-equilibration to 2.3 % B in 1.05 min at a flow rate of 0.85 mL min^−1^. The column (Acquity BEH Shield RP18 column, 50 mm x 2.1 mm, 1.7 µm, Waters) was maintained at 45 °C and sample temperature was kept constant at 14 °C. Monobromobimane conjugates were detected by fluorescence at 480 nm after excitation at 380 nm and quantified using ultrapure standards (Sigma).

For determination of intracellular BCKA content, the same acidic extracts were used. For derivatization with DMB (1,2-diamino-4,5-methylendioxybenzene), 30 µL extract were mixed with 30 µL DMB derivatization reagent (5 mM DMB, 20 mM sodium hydrosulfite, 1 M 2-mercaptoethanol, 1.2 M HCl) and incubated at 100 °C for 45 min. After 10 min centrifugation, the reaction was diluted with 240 µL 10 % acetonitrile. UPLC system, column and solvent were used as described above. Baseline separation of DMB derivates was achieved by increasing the concentration of acetonitrile (B) in buffer A as follows: 2 min 2 % B, 4.5 min 15 % B, 10.5 min 38 % B, 10.6 min 90 % B, hold for 2 min and return to 2 % B in 3.5 min.

The separated derivates were detected by fluorescence (Acquity FLR detector, Waters, OPD: excitation: 350 nm, emission: 410 nm; DMB: excitation: 367 nm, emission: 446 nm) and quantified using ultrapure standards (Sigma). Data acquisition and processing were performed with the Empower3 software suite (Waters).

### Oxygen consumption measurements

Respiration of roots of 16-day-old seedlings grown on ½ MS vertical agar plates in standard long-day condition was analysed using a Clark-type oxygen electrode (Oxygraph, Hansatech) as described in Wagner et al. (2015). Roots from 15-30 seedlings (about 100 mg fresh weight) were cut with a scalpel, rolled and plugged in the oxygraph chamber filled with 1.2 mL incubation medium (5 mM KCl, 10 mM MES, and 10 mM CaCl_2_, pH 5.8). After recording the basal root respiration (5 min) oxygen consumption was followed in response to the sequential addition of the inhibitors 25 µL of KCN (potassium cyanide) at final concentration of 4 mM and 25 μL of pGal (propyl gallate) at final concentration of 200 μM. The measure was repeated switching the order of the inhibitors. The results were normalized on the dry weight of the roots (DW) and on the base respiration rate without inhibitors.

### Gene expression analysis

8-day-old seedlings growing on agar plates in long-day condition were collected and flash-frozen in liquid nitrogen before powdering with metal beads using a TissueLyser II (Qiagen). Total RNA was isolated using the NucleoSpin® RNA Kit (Thermo) and cDNA was synthetized from 10 µg RNA using RevertAid First strand cDNA Synthesis Kit (Thermo) following the supplier instructions. qRT-PCR was performed with PerfeCTa SYBR Green FastMix (Quanta) in a 384-well plate using a CFX96 Real-Time PCR Detection System. The two reference genes *SAND* and *TIP41* were used for normalization. Three technical replicates for each of three independent biological replicates were performed for each experiment. The primers used are listed in Supplemental Table S6. PCR efficiency for each primer pair was assessed by conducting calibration dilution curves in dedicated run.

### Statistical analyses and data plotting

Quantified data were plotted and statistically analysed with GraphPad Prism 6. All the box plots show the median (center line) that divides the first to the third quartile, min and max values are indicated by whiskers, points represent individual values of the whole data set. Outliers were automatically identified with ROUT Method (Q = 10 %) and not shown. Significant differences were calculated according to Student’s *t*-test (two-tailed with 95 % confidence interval) or one-way ANOVA with Tukey’s multiple comparisons (with 95 % confidence interval). Asterisks represent significant differences (*=P ≤ 0.1, **= P ≤ 0.01, ***= P ≤ 0.001, ****= P ≤ 0.0001, ns: not significant).

## Accession numbers

*GRXS15* (AT3G15660), *ACO3* (AT2G05710), *ACO2* (AT4G26970), *LIP1* (AT2G20860), *OAS-TL C* (AT3G59760), *SERAT2;2* (AT3G13110), *H protein 1* (AT2G35370), *H protein 2* (AT2G35120), *H protein 3* (AT1G32470), *PDC-E2 1* (AT3G52200), *PDC-E2 2* (AT3G13930), *PDC-E2 3* (AT1G54220), *PDC-E2 4* (AT3G25860), *OGDC-E2 1* (AT5G55070), *SAND* (AT2G28390), *TIP41* (AT4G34270).

## Acknowledgements

We thank Timon Könen, Lisa Zander and Juan Carlos Davila Frantzen for support in plant maintenance and basic phenotype characterization. We also thank Nicolas Rouhier and Jonathan Przybyla-Toscano for providing the LIP1 antibody.

