## Supplemental Data for "Altered iron-sulfur cluster transfer in Arabidopsis mitochondria reveals lipoyl synthase as a Janus-faced enzyme that generates toxic sulfide"

Supplemental Information includes 20 Figures and 6 Tables.

**A**

| Line | Full line name | Genetic background | Reference |
|---|---|---|---|
| WT | Wild type | Col-0 | |
| *grxs15-1* | *grxs15-1* | SALK_112767 | Ströher *et al.*, 2016; Moseler *et al.*, 2021 |
| *amiR* | GRXS15$^{amiR}$ | Col 0 | Srtöher et *al.*, 2016 |
| *K83A #3* | *grxs15-3 UBQ10pro:GRXS15 K83A #3* | GK-837C05 | Moseler et *al.*, 2015 |
| *K83A #4* | *grxs15-3 UBQ10pro:GRXS15 K83A #4* | GK-837C05 | Moseler *et al.*, 2015 |

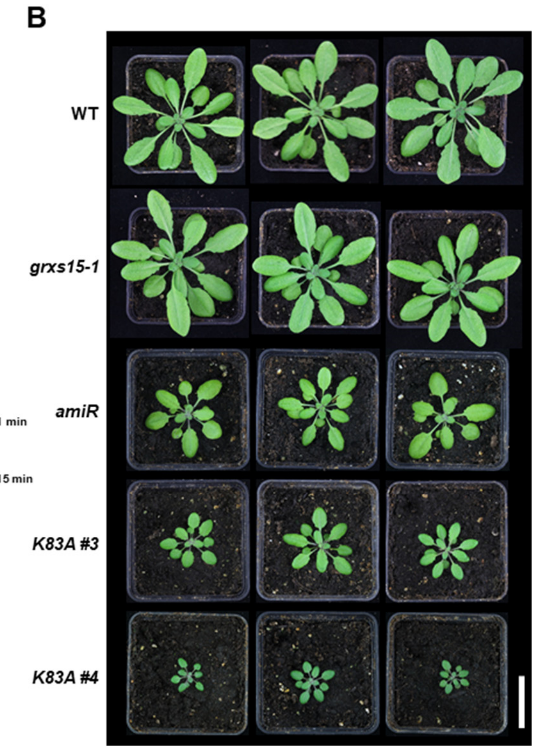


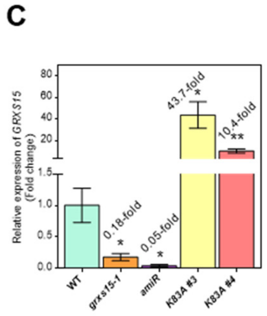


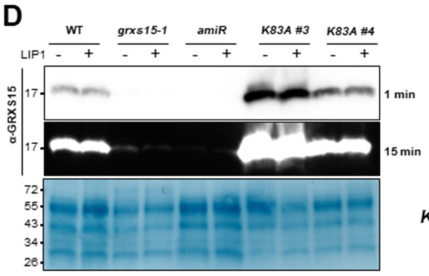


**Supplemental Figure S1. Phenotypes of different *grxs15* mutants. A,** The mutant collection includes two knock-down lines, *grxs15-1* and GRXS15$^{amiR}$ (*amiR*), with decreased amounts of *GRXS15* transcript of about 18 % and 5 %, respectively. In a complementary approach, GRXS15 K83A protein variants were used to rescue the embryo lethal *grxs15-3* (GK-837C05) null mutant in line #*3* and #*4* are driven by a strong ubiquitous promoter (Moseler et al., 2015). **B,** Representative photos of 5-week-old plants grown on soil under long-day conditions. Scale bar: 3 cm. **C**, Relative expression level of *GRXS15* transcripts determined by qRT-PCR in wild type (WT) and the four *grxs15* mutants shown in panel b. The presented data are means ± SEM ($n$ = 3). Student's $t$-test (α = 0.05) has been performed for all mutants (pairwise comparison to wild type) and significant differences are indicated at P < 0.1 (*) and P < 0.01 (**). Changes in comparison to expression in the wild type are shown in fold-changes. *P*-values: Supplemental Data Set 6. **D,** Protein gel blot analysis with primary antibodies raised against GRXS15. 15 µg of mitochondria isolated from seedlings grown in hydroponic culture were used. The blot is shown with two exposure times. Exposure for 15 minutes allows detection of minute amounts of GRXS15 in *grxs15-1* and *amiR*. Amido Black staining of the PVDF membrane serves as loading control. The plant phenotypes (panel b) and the blot exposed for 1 min are also shown in Figure 3 of the main manuscript.

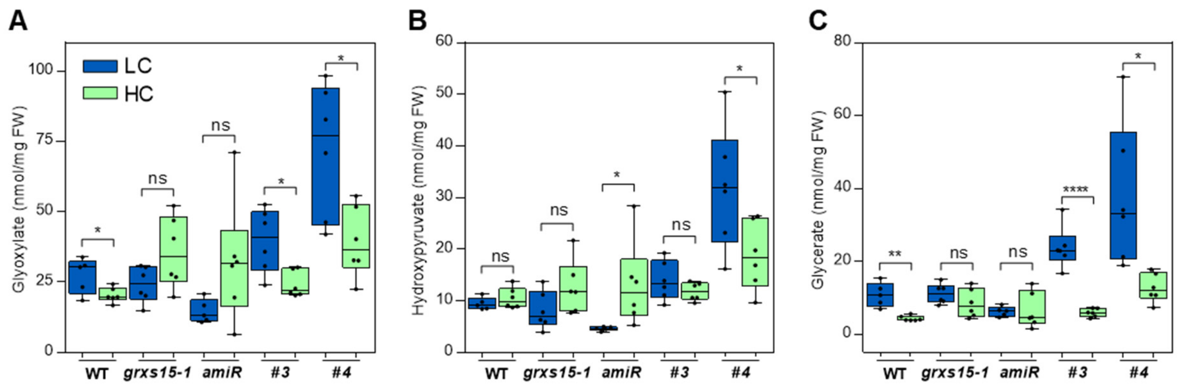


**Supplemental Figure S2. Suppression of photorespiration has an impact on photorespiratory intermediates glyoxylate, hydroxypyruvate and glycerate in mutants complemented with GRXS15 K83A.** Metabolic amounts of 8-week-old plants of hydroxypyruvate (**A**), glycerate (**B**) and glyoxylate (**C**) in wild type and *grxs15* mutants *grxs15-1*, *amiR*, *K83A #3* and *#4* grown in low carbon dioxide condition (LC: 390 ppm $CO_2$) and high carbon dioxide condition (HC: 5,000 pm $CO_2$). Box plots show the median as centre line with the box for the first to the third quartile and whiskers indicating min and max values of the whole dataset (*n* = 5-6). Asterisks represent significant differences (*=P ≤ 0.1, ****= P ≤ 0.0001, ns: not significant) calculated according to Student's *t*-test (α = 0.05). *P*-values: Supplemental Data Set 7.

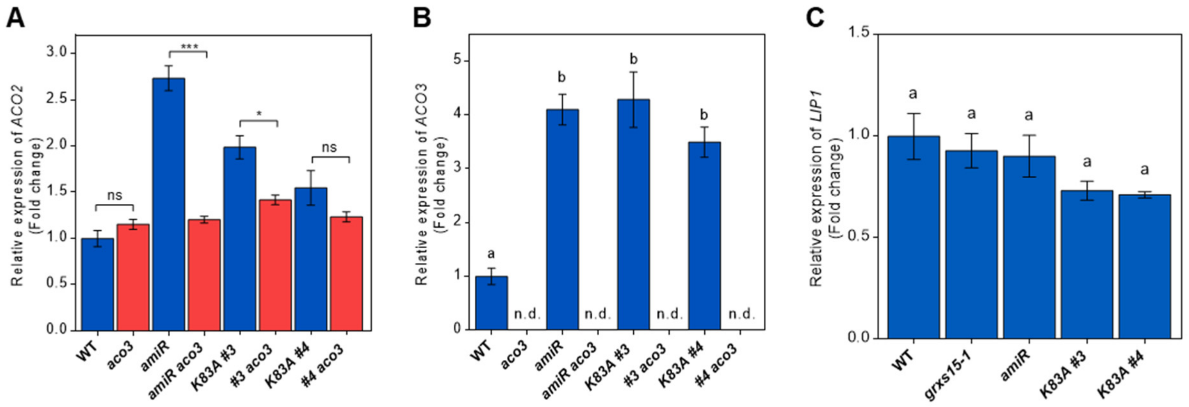


**Supplemental Figure S3. Expression level of *ACO2*, *ACO3* and *LIP1* in the *grxs15* mutants.** Relative expression level of *ACO2* (**A**), *ACO3* (**B**) and *LIP1* (**C**) transcripts determined by qRT-PCR in wild type and the four *grxs15* mutants. The presented data are means ± SEM ($n$ = 3). Student's $t$-test has been performed for all mutants against the wild type ($\alpha = 0.05$), different letters indicate significant statistically different groups and asterisks indicate $P < 0.1$ (*), $P < 0.001$ (***), ns: not significant and n.d.: not detected. *P*-values: Supplemental Data Set 8.

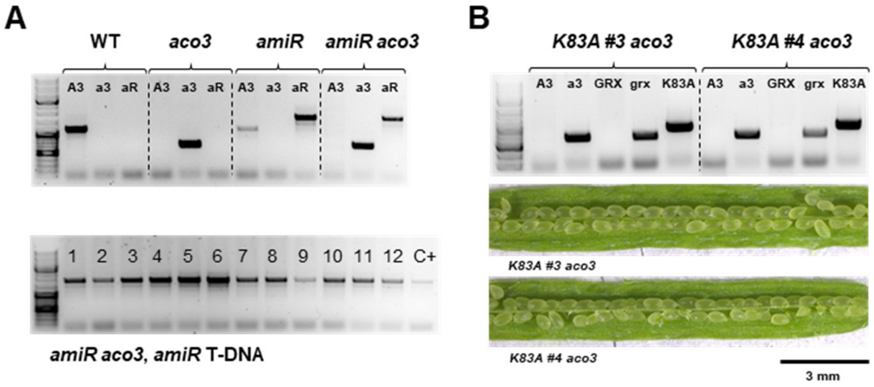


**Supplemental Figure S4. Confirmation of genotypes for crosses of *aco3* and *grxs15.* A,** Genotyping of the cross *amiR* × *aco3*. A3: *ACO3* WT allele, a3: *aco3* T-DNA; aR: *amiR* T-DNA. The panel at the bottom shows PCR results for 12 independent plants to confirm the selected line *amiR* × *aco3* as homozygous for *amiR*. **B,** Genotyping of the crosses *K83A #3* and *#4* × *aco3*. GRX: *GRXS15* WT allele; K83A: T-DNA carrying the lysine mutation. The panels at the bottom show opened siliques to confirm the absence of aborted seeds.

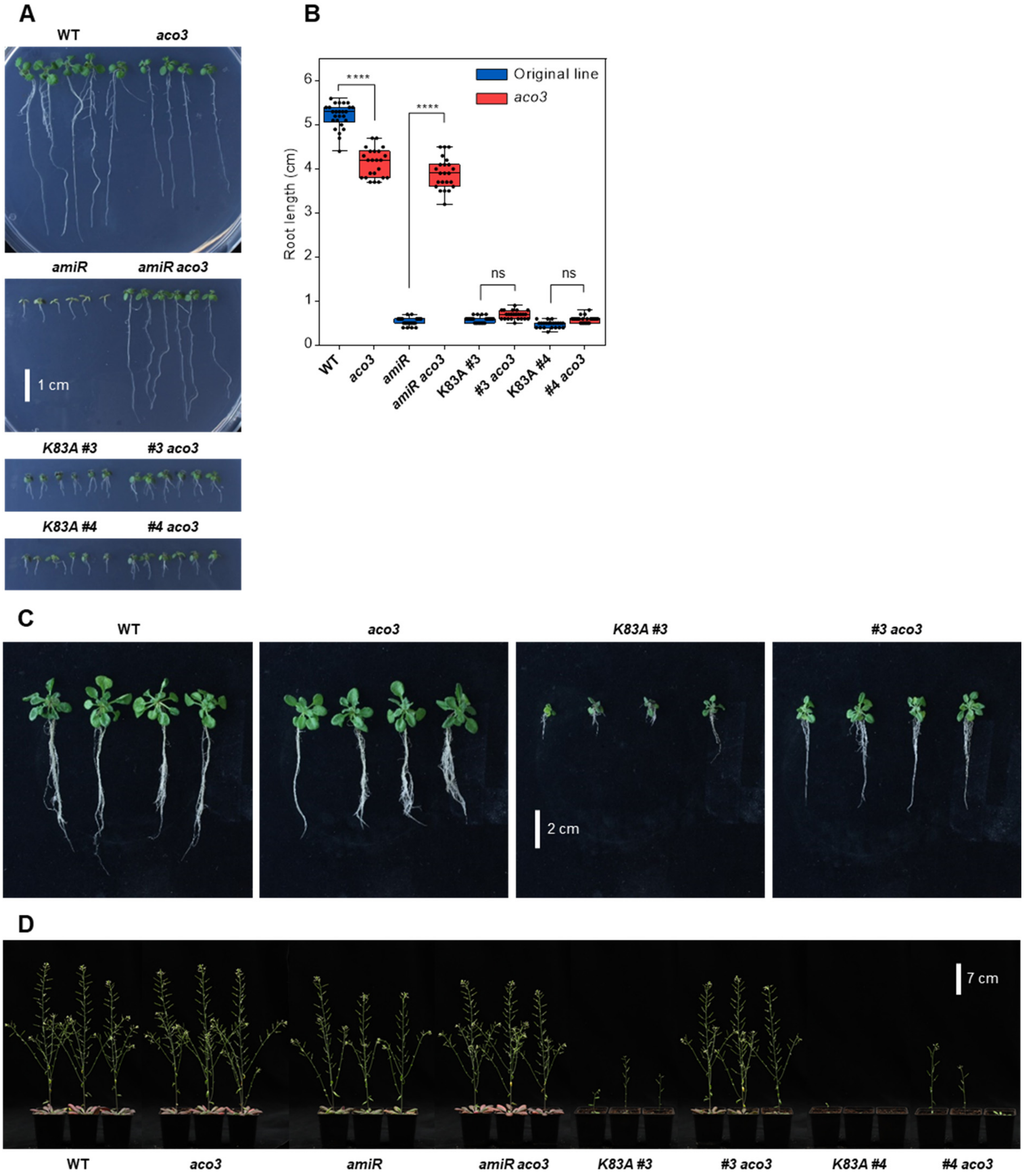


**Supplemental Figure S5. Loss of *ACO3* partially suppressed the dwarfism in *grxs15* mutants. A,** 10-day-old seedlings grown on ½ MS vertical agar plates under long-day conditions. **B,** Root length analysis ($n$ = 22-26). Data for WT and the different *grxs15* mutants are shown in blue, data for *aco3* and the respective crosses are shown in red. Asterisks indicate P < 0.0001 (****) and ns: not significant, calculated according to Student's *t*-test (α = 0.05). *P*-values: Supplemental Data Set 9. **C,** 17-day-old seedlings grown on vertical agar plates under long-day conditions. **D,** Photo of three representative 8-week-old plants of *aco3* and crosses compared with their respective parental lines. All plants were grown on soil under long-day conditions.

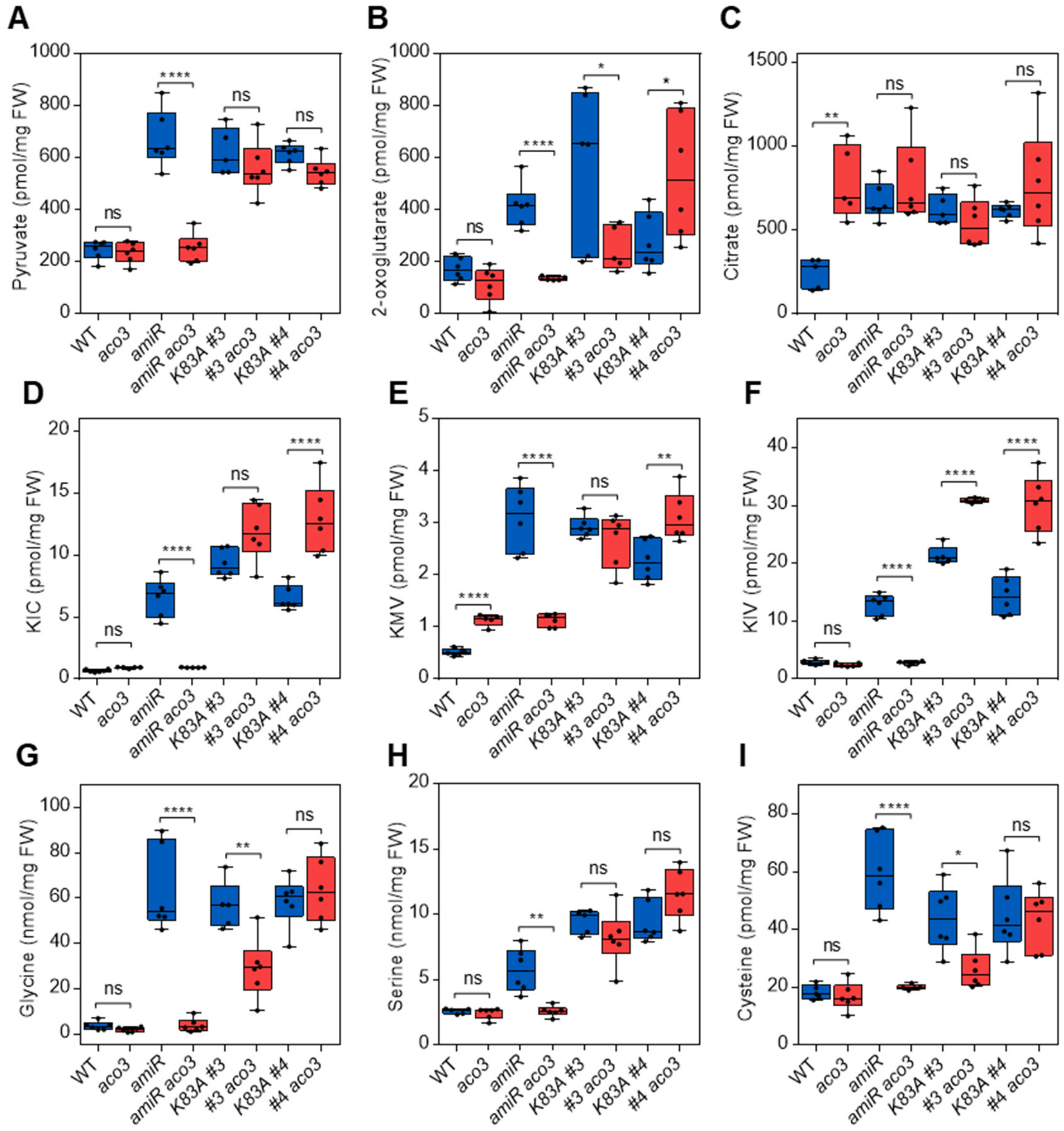


**Supplemental Figure S6. Loss of *ACO3* has an impact on the metabolite signature.** Metabolite signatures of 8-day-old seedlings. Analysis of TCA cycle metabolites pyruvate (**A**) and 2-OG (**B**), which are substrates of PDC and OGDC, respectively, and citrate (**C**), which is converted by aconitases; three branched chain α-keto acids (KIC, KMV and KIV) (**D-F**), which are all degraded by BCDHC, and the amino acids glycine (**G**) and serine (**H**) that are linked to photorespiration, and the activity of the mitochondrial glycine dehydrogenase complex (GDC). Cysteine (**i**) is synthesized from its immediate precursor serine. All box plots show the median as centre line with the box for the first to the third quartile and whiskers indicating min and max values of the whole dataset ($n$ = 5-6). Asterisks represent significant differences (*=$P \leq 0.1$, **= $P \leq 0.01$ ***= $P \leq 0.001$ ****= $P \leq 0.0001$, ns: not significant) calculated according to one-way ANOVA with Tukey's multiple comparisons test (α = 0.05). Data for WT and the different *grxs15* mutants are shown in blue, data for *aco3* and the respective crosses are shown in red. *P*-values: Supplemental Data Set 10.

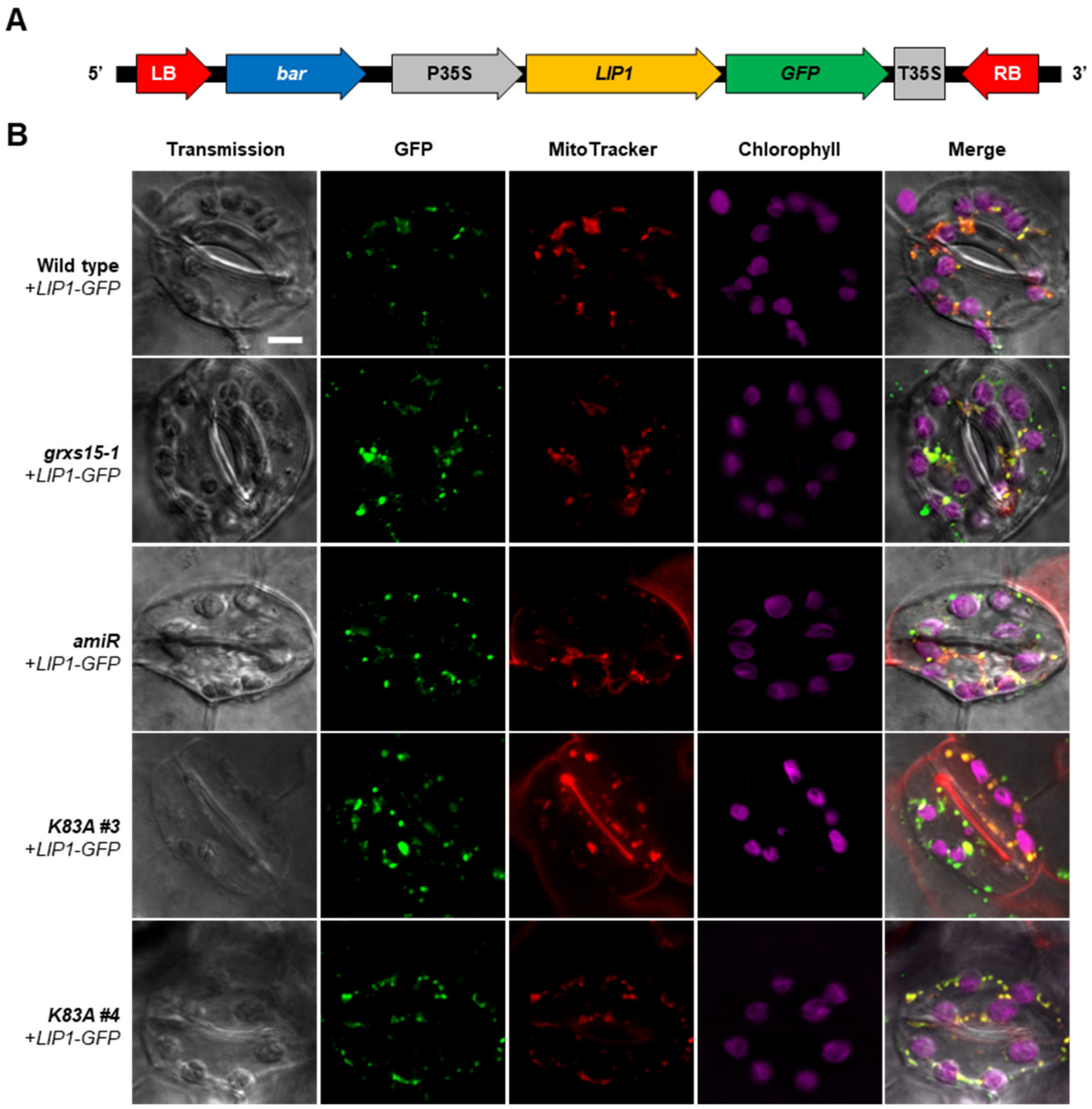


**Supplemental Figure S7. Transformed plants expressed LIP1-GFP in mitochondria. A,** Representation of the construct used for the overexpression of LIP1 with a C-terminal GFP for the subcellular localization. The gene coding for *LIP1* includes the original signal peptide for mitochondrial targeting. **B,** Confocal microscopy images show stomata from 7-day-old seedlings stably expressing LIP1-GFP in different genetic backgrounds (wild type, *grxs15-1*, *amiR*, *K83A #3* and *#4*). The figure shows transmission images, GFP fluorescence ($\lambda_{ex}$ = 488 nm; $\lambda_{em}$ = 520 nm), MitoTracker Orange staining for mitochondria ($\lambda_{ex}$ = 543 nm; $\lambda_{em}$ = 597 nm), chlorophyll autofluorescence ($\lambda_{ex}$ = 633 nm; $\lambda_{em}$ = 675 nm) and the merged images. For each line one representative set of images selected from three independent transgenic lines is shown. Scale bar: 5 µm.

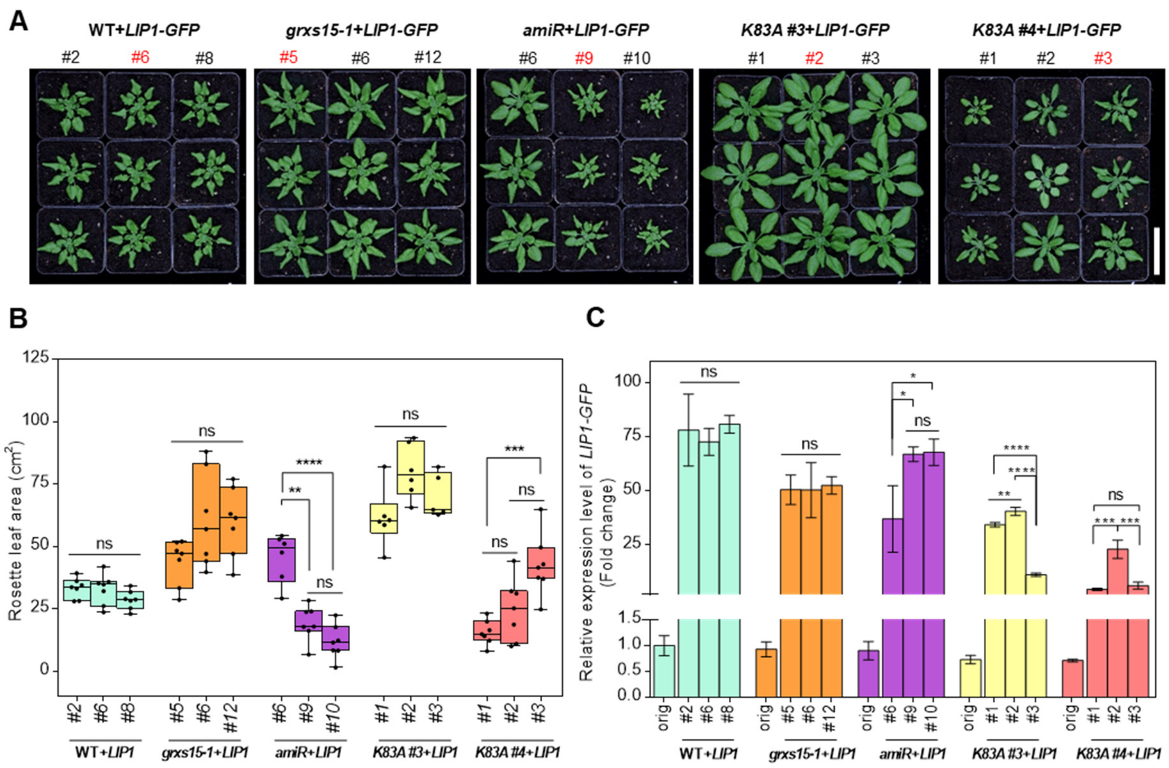


**Supplemental Figure S8. Overexpression of *LIP1-GFP* results in phenotypic effects that depend on the respective genetic background and vary in several independent lines. A,** 5-week-old plants of three independent stable lines transformed with pSS01_*35S$_{pro}$*:*LIP1-GFP* grown under long-day conditions (wild type+*LIP1-GFP*: numbers #2, #6, #8; *grxs15-1+LIP1-GFP*: #5, #6, #12; *amiR+LIP1-GFP*: #6, #9, #10; *K83A #3+LIP1-GFP*: #1, #2, #3, and *K83A #4+LIP1-GFP*: #1, #2, #3). *LIP1* overexpression lines chosen for all further phenotypic characterization, metabolite analyses and transformation with *OAS-TL C* are indicated with red numbers. Scale bar: 5 cm. **B,** Rosette area of the three independent stable lines after four weeks on soil under log-day conditions ($n$ = 8). Box plots show the median as center line with the box for the first to the third quartile and whiskers indicating min and max values of the whole dataset. **C,** Relative expression level of *LIP1* transcripts determined by qRT-PCR in wild type and the four *grxs15* mutants. RNA was isolated from 10-day-old seedlings. The presented data are means ± SEM ($n$ = 3). Different letters indicate significantly different groups calculated according to one-way ANOVA with Tukey's multiple comparisons test separately for each genotype plus the respective genetic background (orig) ($\alpha$ = 0.05). *P*-values: Supplemental Data Set 11.

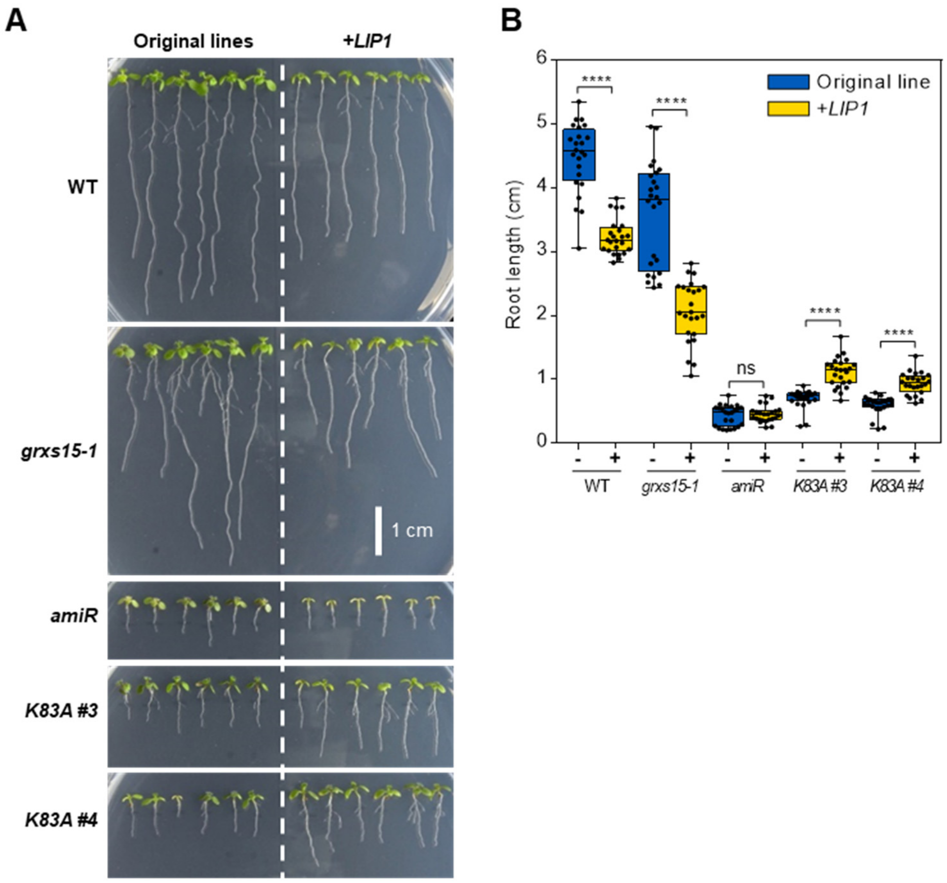


**Supplemental Figure S9. *LIP1* overexpression affects root length. A,** 10-day-old seedlings grown on ½ MS vertical agar plates in long-day conditions. **B,** Analysis of root length of *LIP1-GFP* overexpression lines compared with their respective genetic background. The box plot shows the median as centre line with the box for the first to the third quartile and whiskers indicating min and max values of the whole dataset (*n* = 24). Asterisks indicate P < 0.0001 (****) and ns: not significant, calculated according to Student's *t*-test (α = 0.05). *P*-values: Supplemental Data Set 12.

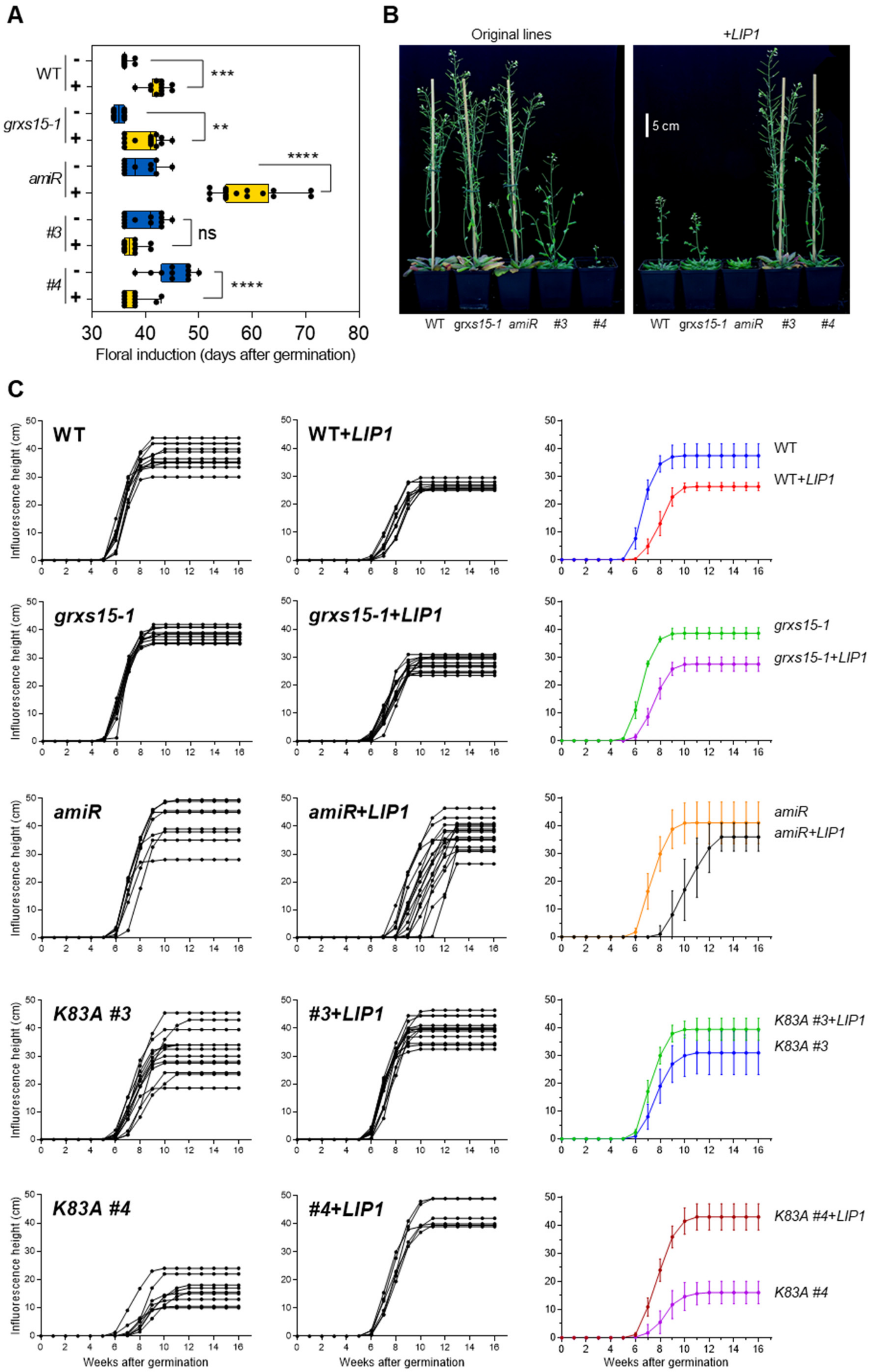
A
B
C
WT
grxs15-1
amiR
#3
#4
***
**
****
ns
****
Floral induction (days after germination)
Original lines
+LIP1
5 cm
WT grxs15-1 amiR #3 #4
WT grxs15-1 amiR #3 #4
Influorescence height (cm)
Weeks after germination
WT
WT+LIP1
grxs15-1
grxs15-1+LIP1
amiR
amiR+LIP1
K83A #3
#3+LIP1
K83A #3+LIP1
K83A #4
#4+LIP1
K83A #4+LIP1

**Supplemental Figure S10. *LIP1* overexpression affects floral induction and inflorescence development. A,** Floral induction. The data show the time after germination when the developing flower bud became visible. The box plot shows the median as centre line with the box for the first to the third quartile and whiskers indicating min and max values of the whole dataset (*n* = 11-17). Asterisks represent significant differences (**= P ≤ 0.01, ***= P ≤ 0.001, ****= P ≤ 0.0001, ns: not significant) calculated according to one-way ANOVA with Tukey's multiple comparisons test (α = 0.05). Data for WT and the different *grxs15* mutants are shown in blue, data for the respective lines overexpressing *LIP1* are shown in yellow. **B** and **C,** Documentation of inflorescence development. (**B**) Flower stalk height nine weeks after germination for non-transformed control plants (left) and plants overexpressing *LIP1-GFP* (right). All plants were grown on soil under long-day conditions. (**C**) Development of the main inflorescence. Plants were grown continuously under long-day conditions and measured three times per week. Panels on the right show the average inflorescence height ± SD calculated from data for individual plants shown on the left (*n* = 6-17). *P*-values: Supplemental Data Set 13.

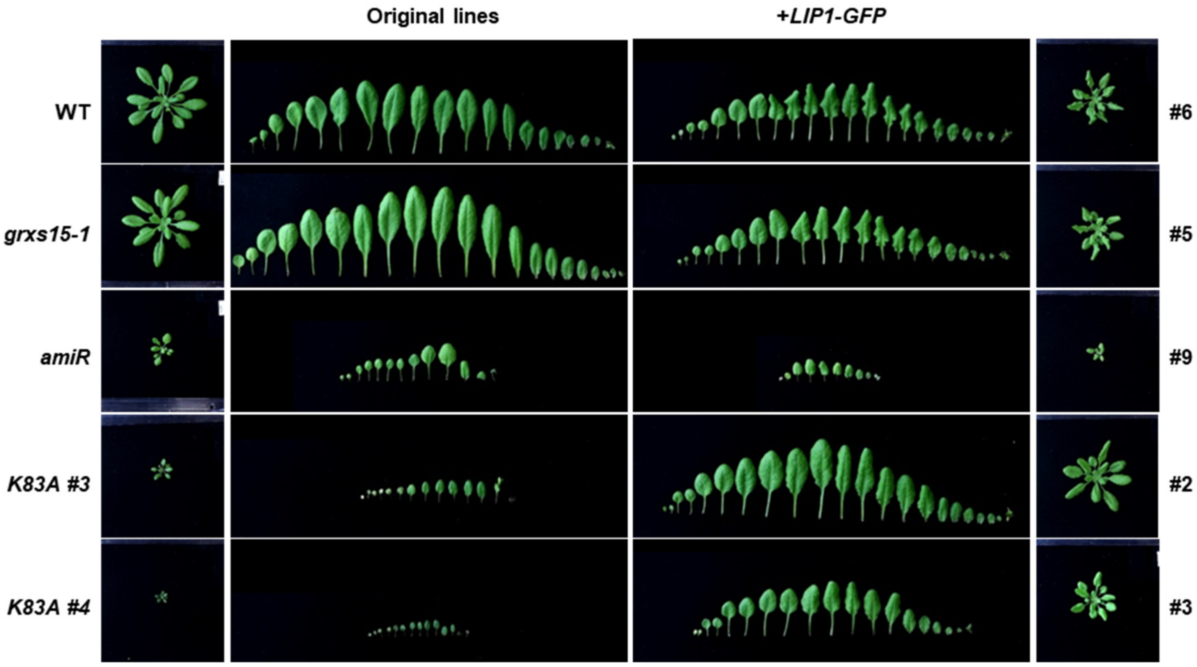


**Supplemental Figure S11. *LIP1* overexpression induced a curly leaf phenotype.** Plants stably overexpressing *LIP1-GFP* were grown on soil under long-day conditions for 32 days. At the time of harvest, the root system was removed and rosettes were dissected to isolate all leaves according to their developmental age from the older leaves on the left to the youngest leaves on the right. The numbers on the right identify the selected lines (see Supplemental Figure S8A).

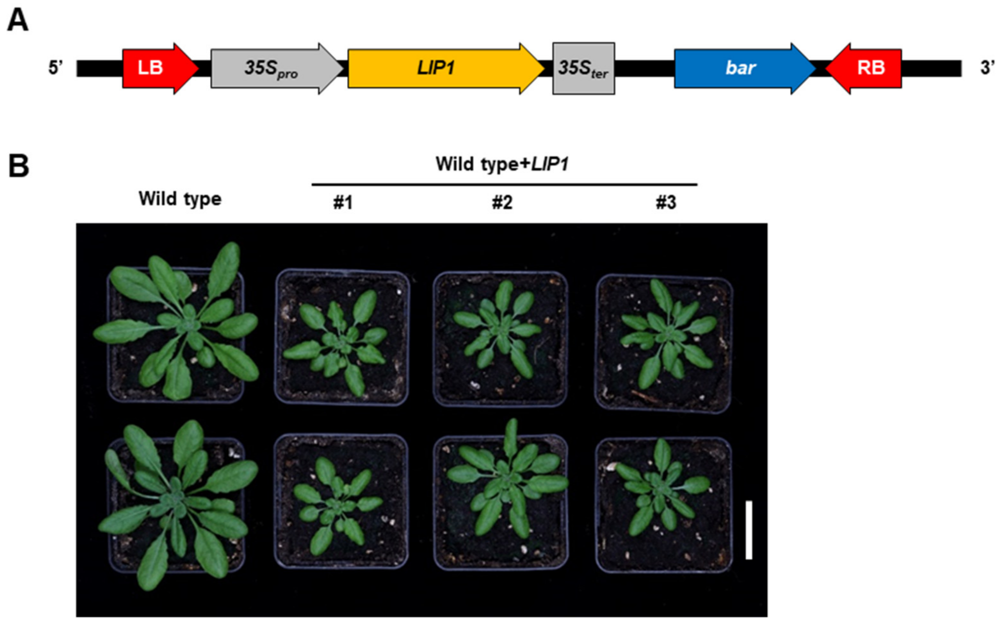


**Supplemental Figure S12. Overexpression of *LIP1* without a GFP tag is deleterious. A,** Schematic representation of the construct used for the overexpression of *LIP1* without GFP. The T-DNA is 3,890 bp long and uses the vector pB7WG2 as backbone (Karimi et al., 2002). **B,** Representative images of three independent stable lines transformed with pB7WG2_$35S_{pro}$:*LIP1* grown for five weeks under long-day conditions. Scale bar: 3 cm.

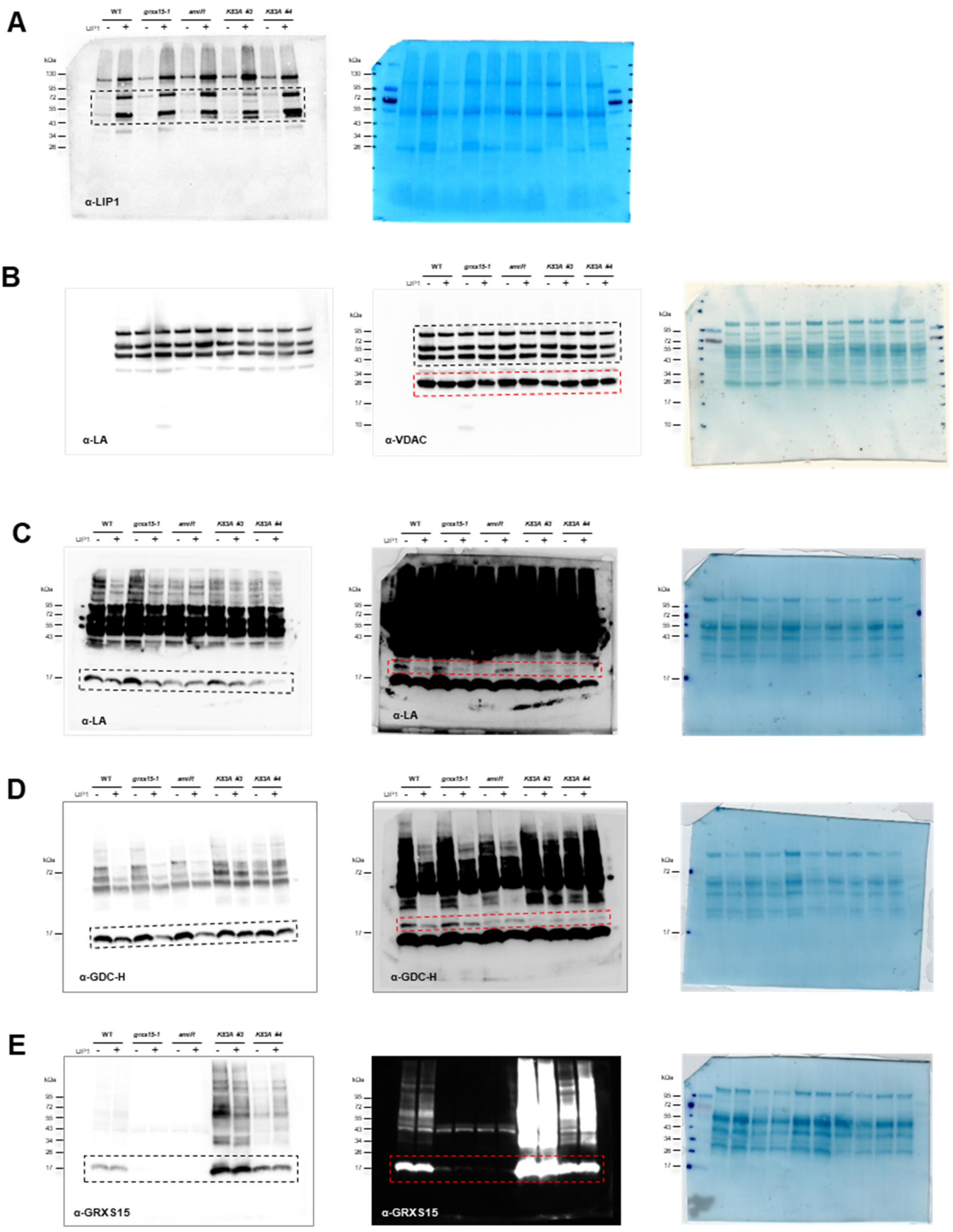


**Supplemental Figure S13. Uncropped western blot images. A,** 15 µg of proteins extract from mitochondria isolated from plants grown on soil were transferred with semidry blotting on PVDF membrane which was incubated with a 1:1,000 dilution of primary antibody raised against LIP1 (Moseler et al., 2021) for 2 h at room temperature. The dashed rectangle delineates the area shown in Figure 3G. **B,** 15 µg of protein extracted from mitochondria isolated from seedlings grown in hydroponic were transferred with semidry blotting on PVDF membrane which was incubated with a 1:1,000 dilution of primary antibody raised against lipoylated proteins (Abcam, ab58724) over-night at 4 °C. The same membrane was re-incubated with 1:5,000 dilution of primary antibody raised against VDAC1-5 HRP-conjugated (Agrisera, AS07 201-HRP). The dotted rectangles delineate the areas for lipoylated proteins (grey) and VDAC (red) shown in Figure 3G. **C,** 20 µg of protein extracted from mitochondria isolated from seedlings grown in hydroponics were transferred by wet blotting onto a PVDF membrane, which was incubated with a 1:500 dilution of primary antibody raised against lipoylated proteins for 3 h at room temperature. The dotted rectangles delineate the areas shown in Figure 3G. The grey line shows the lipoylated GDC-H isoforms 1 and 3 and in the red line part of the same blot after prolonged exposure time to highlight the isoform GDC-H2. **D,** Similar to c, 20 µg of protein extracted from mitochondria isolated from seedlings grown in hydroponic were transferred with wet blotting on PVDF membrane which was incubated with a 1:5,000 dilution primary antibody raised against GDC-H proteins (Agrisera As05 074) for 3 h at room temperature. The grey dashed rectangle highlights GDC-H isoforms 1 and 3 and the red rectangle a prolonged exposure time to highlighted the isoform GDC-H2. Both areas are shown in Figure 3G. **E,** 20 µg of protein extracted from mitochondria isolated from seedlings grown in hydroponic were transferred with wet blotting on PVDF membrane which was incubated with a 1:2,500 dilution of primary antibody raised against GRXS15 (Moseler et al., 2015) for 2 h at room temperature. The dotted rectangle delineates the area shown in Figure 3G and Supplemental Figure S1. Highlighted in red is the same part of the blot with an extended exposure time as displayed in Supplemental Figure S1.

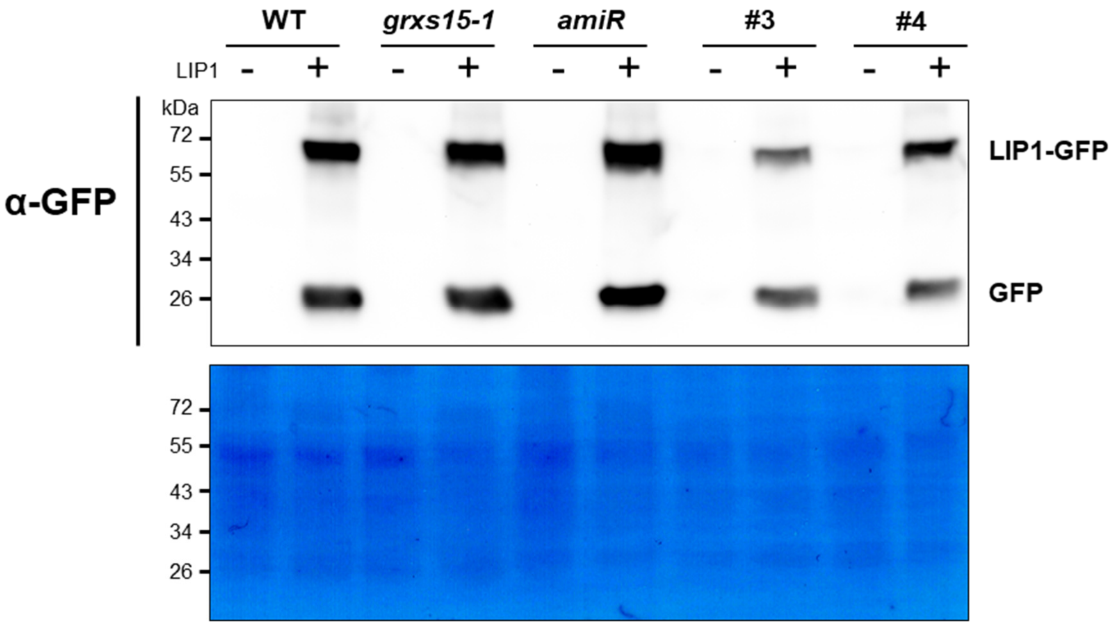


**Supplemental Figure S14. Protein gel blot analysis with antiserum raised against GFP.** Protein gel blot analysis with primary antibodies raised against GFP (Thermo, A-6455). 15 µg of protein extracted from mitochondria isolated from seedlings grown in hydroponic culture were loaded. Amido Black staining of the PVDF membrane serves as loading control. The two bands indicate that a fraction of the LIP1-GFP fusion protein got cleaved.

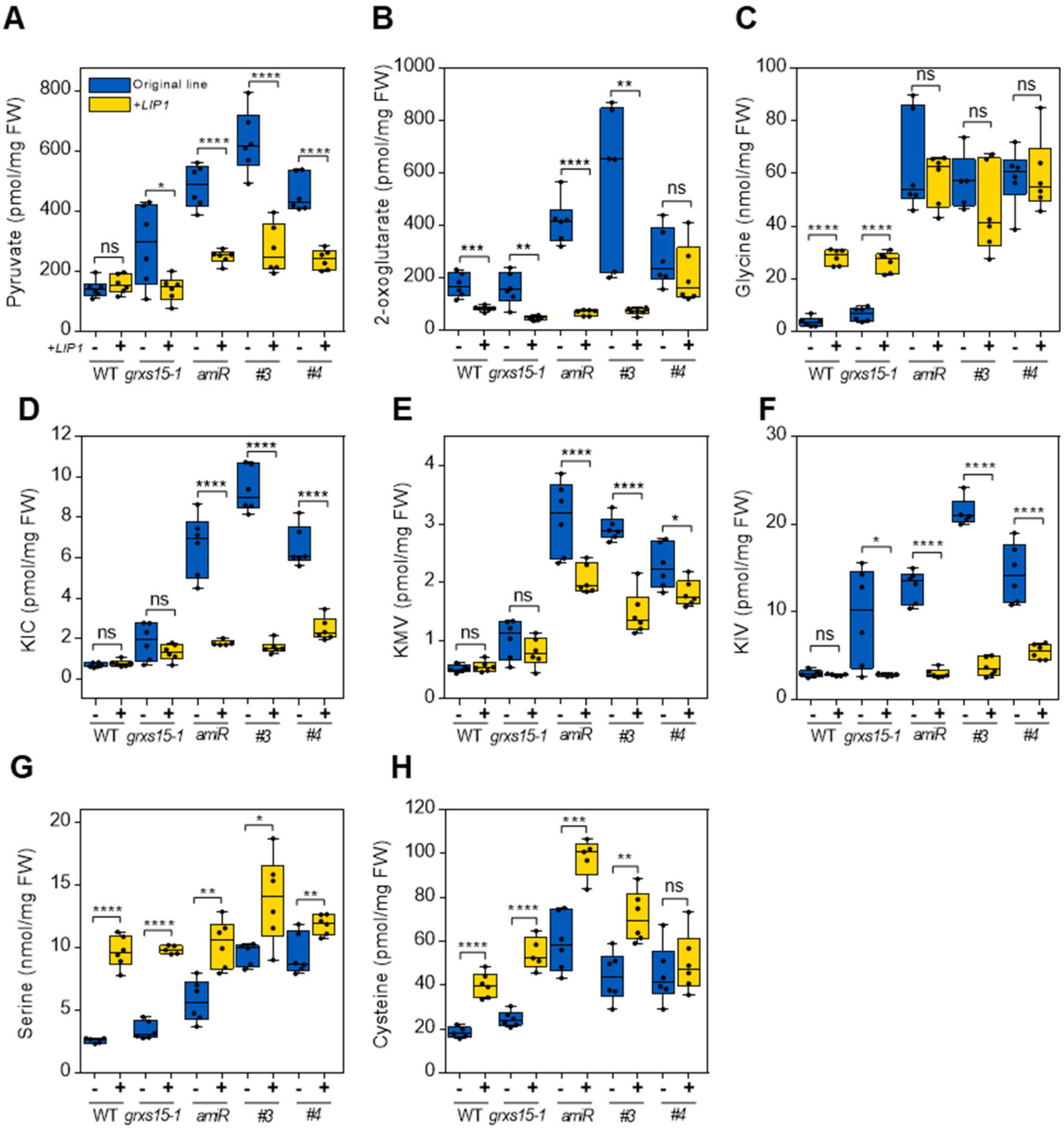


**Supplemental Figure S15. Overexpression of *LIP1* causes pronounced metabolic changes. A-F,** Concentration of substrates of LA-dependent mitochondrial enzymes: pyruvate as the substrate of PDC (**A**), 2-oxoglutarate as the substrate of OGDC (**B**), glycine as the substrate of GDC (**C**), and three branched chain α-keto acids (**D-F**), which are all metabolized by BCDHC. **G** and **H,** Serine and cysteine, which are formally derived in parts from glycine. Metabolites were extracted from 8-day-old seedlings grown under long-day conditions. The box plots show the median as centre line with the box for the first to the third quartile and whiskers indicating min and max values of the whole dataset ($n$ = 5-6). Asterisks represent significant differences (*=$P \leq 0.1$, **= $P \leq 0.01$, ***= $P \leq 0.001$, ****= $P \leq 0.0001$, ns: not significant) calculated according to Student's $t$-test ($\alpha = 0.05$). $P$-values: Supplemental Data Set 14. Data for WT and the different *grxs15* mutants are shown in blue (-), data for the respective lines overexpressing *LIP1* are shown in yellow (+).

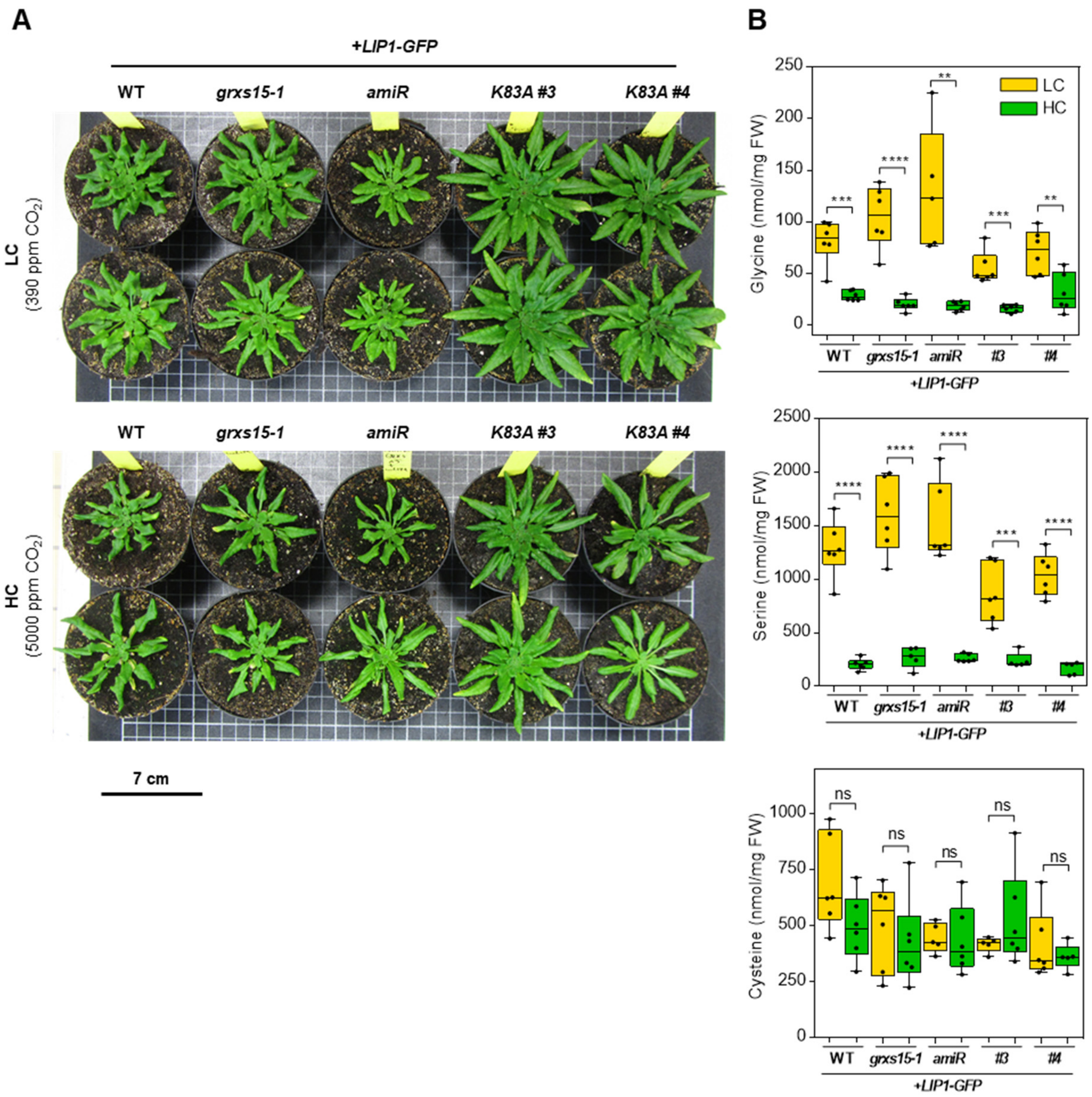


**Supplemental Figure S16. High $CO_2$ has no beneficial effect on *LIP1* overexpression lines. A,** Phenotype comparison between 8-week-old wild type (WT) and *grxs15* mutants stably overexpressing *LIP1-GFP* grown in normal air and high $CO_2$ (390 and 5,000 ppm, respectively) and a 12/12-hour day/night light regime. **B,** Content of glycine, serine and cysteine of 8-week-old plants grown in low (LC, shown in yellow) and high carbon dioxide condition (HC, shown in green) ($n$ = 5-6) (for absolute values see $P$-values: Supplemental Data Set 15). Asterisks represent significant differences (**= $P \leq 0.01$ ***= $P \leq 0.001$ ****= $P \leq 0.0001$, ns: not significant) calculated according to Student's $t$-test (α = 0.05).

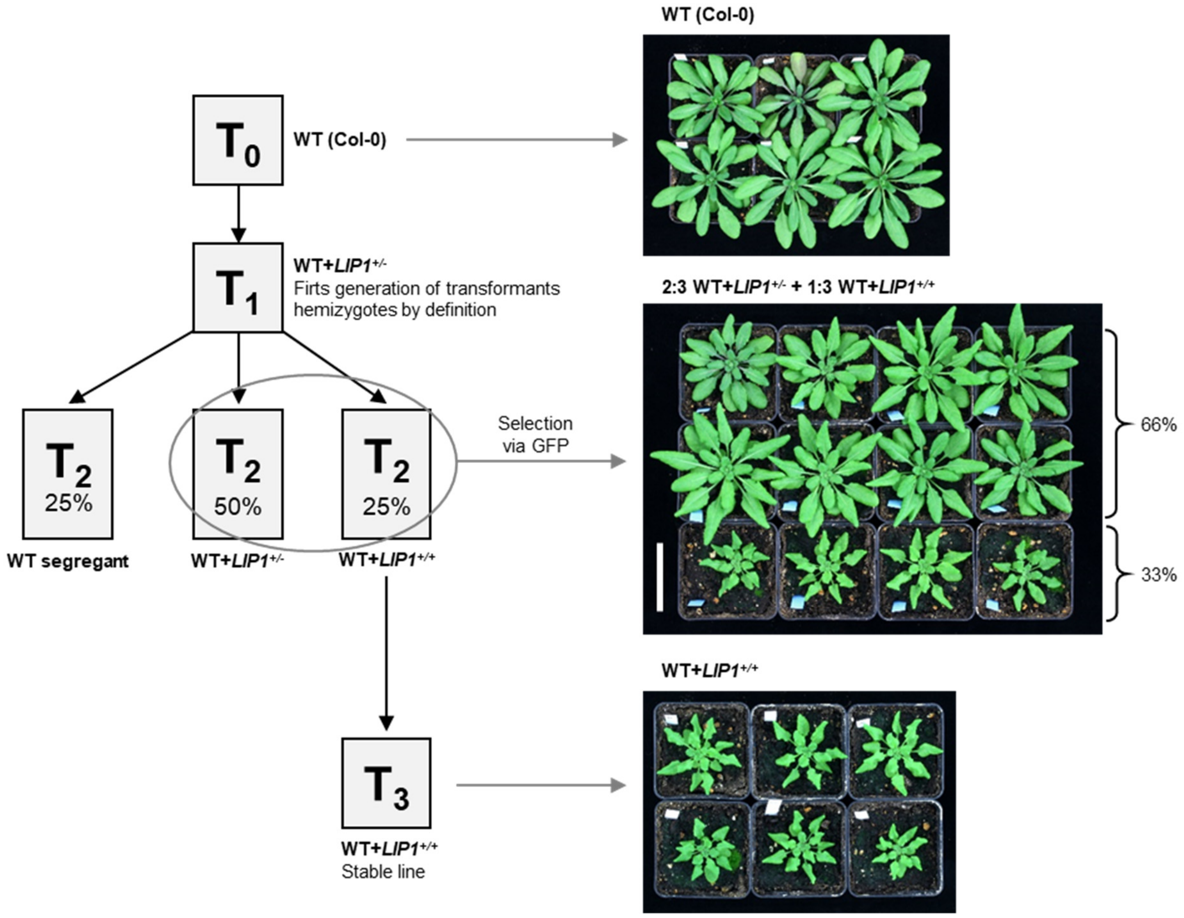


**Supplemental Figure S17. Overexpression of *LIP1* has a gene dosage-dependent effect on wild type (WT) plants.** Representative picture of 5-week-old WT plants overexpressing *LIP1-GFP* in homozygosis (+/+) and hemizygosis (+/-) compared with the original WT. $T_0$ WT plants were transformed for *LIP1-GFP* overexpression, $T_1$ plants were selected via herbicide resistance (BASTA). $T_2$ segregated and transformants were selected by visual inspection for GFP fluorescence. Non-fluorescent WT segregants (25 %) were removed and the remaining fluorescent individuals selected (75 %) subsequently phenotypically separated into two classes presumed to be hemizygous (66 %) and homozygous (33 %) for the *LIP1-GFP* overexpression. Homozygosity was confirmed in $T_3$ populations. Scale bar: 5 cm.

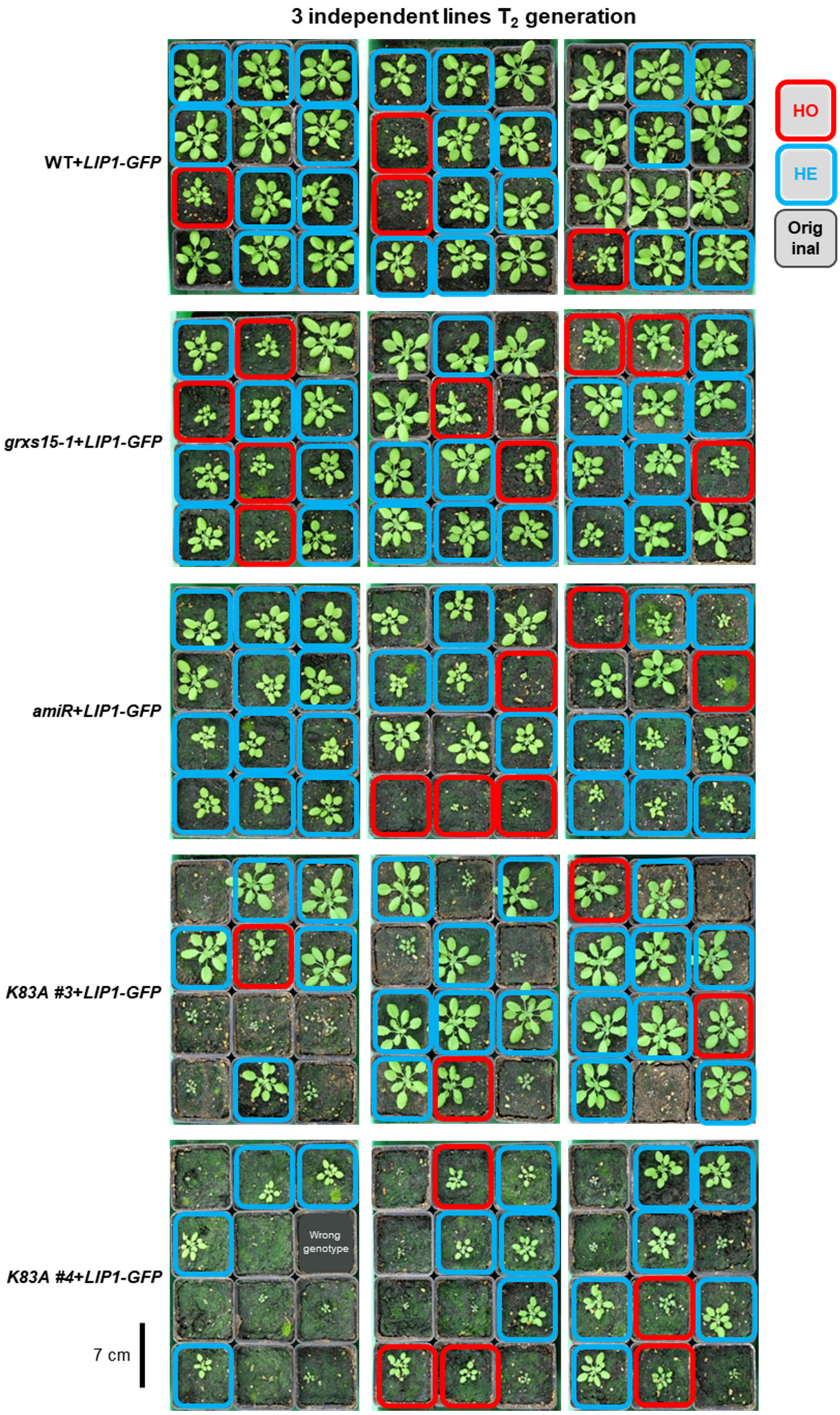
3 independent lines T2 generation
HO
HE
Original
WT+LIP1-GFP
grxs15-1+LIP1-GFP
amiR+LIP1-GFP
K83A #3+LIP1-GFP
K83A #4+LIP1-GFP
Wrong genotype
7 cm

**Supplemental Figure S18. Overexpression of *LIP1* has a gene dosage-dependent effect on WT and GRXS15-deficient mutants.** Photos of 4-week-old plants from the screening of the second generation ($T_2$) for *LIP1-GFP* overexpression. Twelve plants of each independent line selected per genotype were grown on soil in standard long-day conditions. The next generation ($T_3$) was then analysed via GFP to determine the zygosity of the respective mother plant (100 % fluorescent seedlings: homozygotes; 75 % fluorescent seedlings: hemizygotes; 0 % fluorescent seedlings: WT segregants). Scale bar: 7 cm.

A

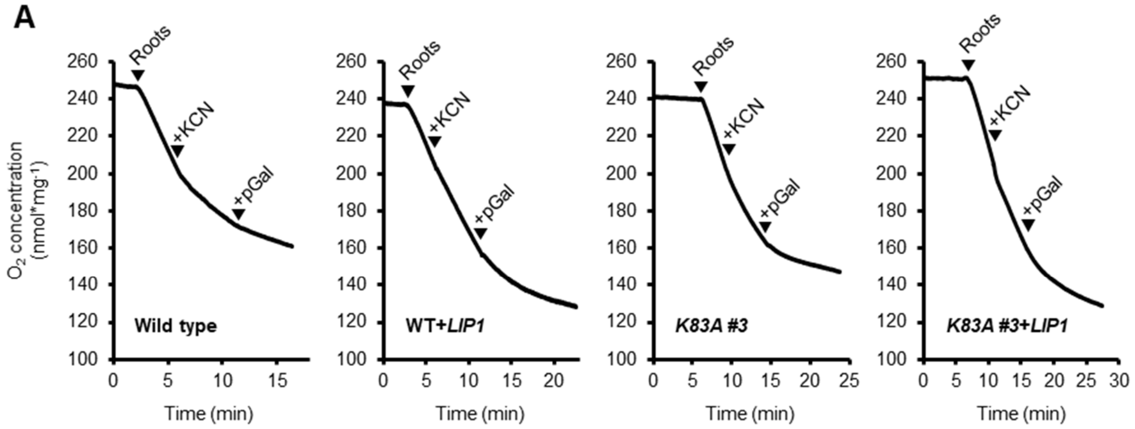


B

| Line | Repl. | DW (mg) | Baseline | Roots | KCN | pGal |
|---|---|---|---|---|---|---|
| WT | #2 | 5.9 | 0.08 | 2.10 | 0.75 | 0.31 |
| | #3 | 4.3 | 0.01 | 1.77 | 0.91 | 0.19 |
| | #4 | 6.3 | 0.10 | 1.35 | 0.64 | 0.15 |
| | #5 | 7.7 | 0.06 | 1.06 | 0.55 | 0.19 |
| WT+*LIP1* | #1 | 6.2 | 0.05 | 3.20 | 2.41 | 0.27 |
| | #2 | 4.8 | 0.08 | 1.92 | 1.77 | 0.19 |
| | #3 | 5.3 | 0.05 | 2.06 | 1.56 | 0.22 |
| | #4 | 10.3 | 0.04 | 2.00 | 1.58 | 0.08 |
| *K83A #3* | #1 | 5.6 | 0.08 | 1.85 | 1.34 | 0.24 |
| | #2 | 5.5 | 0.02 | 2.37 | 1.10 | 0.21 |
| | #3 | 9.2 | 0.04 | 1.79 | 0.73 | 0.07 |
| | #4 | 6.3 | 0.09 | 1.23 | 0.48 | 0.14 |
| | #5 | 5.4 | 0.11 | 2.21 | 0.92 | 0.22 |
| *#3+LIP1* | #1 | 7.2 | 0.04 | 0.90 | 0.58 | 0.10 |
| | #2 | 6.9 | 0.04 | 1.20 | 0.60 | 0.09 |
| | #3 | 8 | 0.03 | 1.59 | 0.94 | 0.18 |
| | #4 | 7.9 | 0.04 | 1.16 | 0.59 | 0.07 |
| | #5 | 6.9 | 0.04 | 1.62 | 1.03 | 0.14 |

C

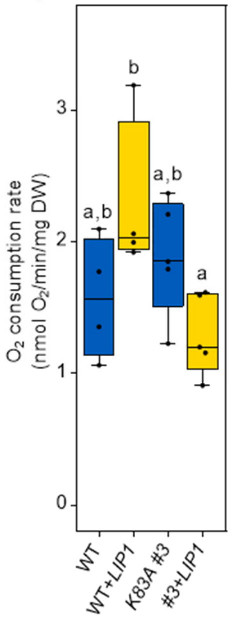


**Supplemental Figure S19. Overexpression of *LIP1* has an impact on respiration of roots. A,** Exemplary respirograms of four genotypes (absolute values highlighted in grey, on table in (**B**). Roots were cut from 16-day-old seedlings grown on ½ MS vertical agar plates and oxygen consumption was followed by time with a Clark-type oxygen electrode. After recording the basal root respiration without inhibitors (baseline), arrows indicate the addition of roots, KCN (potassium cyanide; final concentration 4 mM) for inhibition of COX and pGal (propyl gallate; final concentration 200 µM) to inhibit AOX. After the measurements, the roots were incubated O/N at 60 °C to dry. **B,** Rates of respiration were calculated in a window of 1 minute, directly before the addition of roots or KCN and the end of the measurement after pGal addition by the software Oxygraph Plus (Hansatech Instruments Ltd). The table shows the absolute values of respiration measurements shown in Figure 4E, normalized on the dry weight (DW). For each of the four genotypes four or five replicates were done. **C,** Basic root respiration of all four genotypes (*n* = 4-5). The box plot shows the median as centre line with the box for the first to the third quartile and whiskers indicating min and max values of the whole dataset. Different letters indicate significant differences between treatments calculated according to one-way ANOVA with Tukey's multiple comparisons test (α = 0.05). Supplemental Data Set 16. Data for WT and *K83A #3* are shown in blue, data for the respective lines overexpressing *LIP1* are shown in yellow.

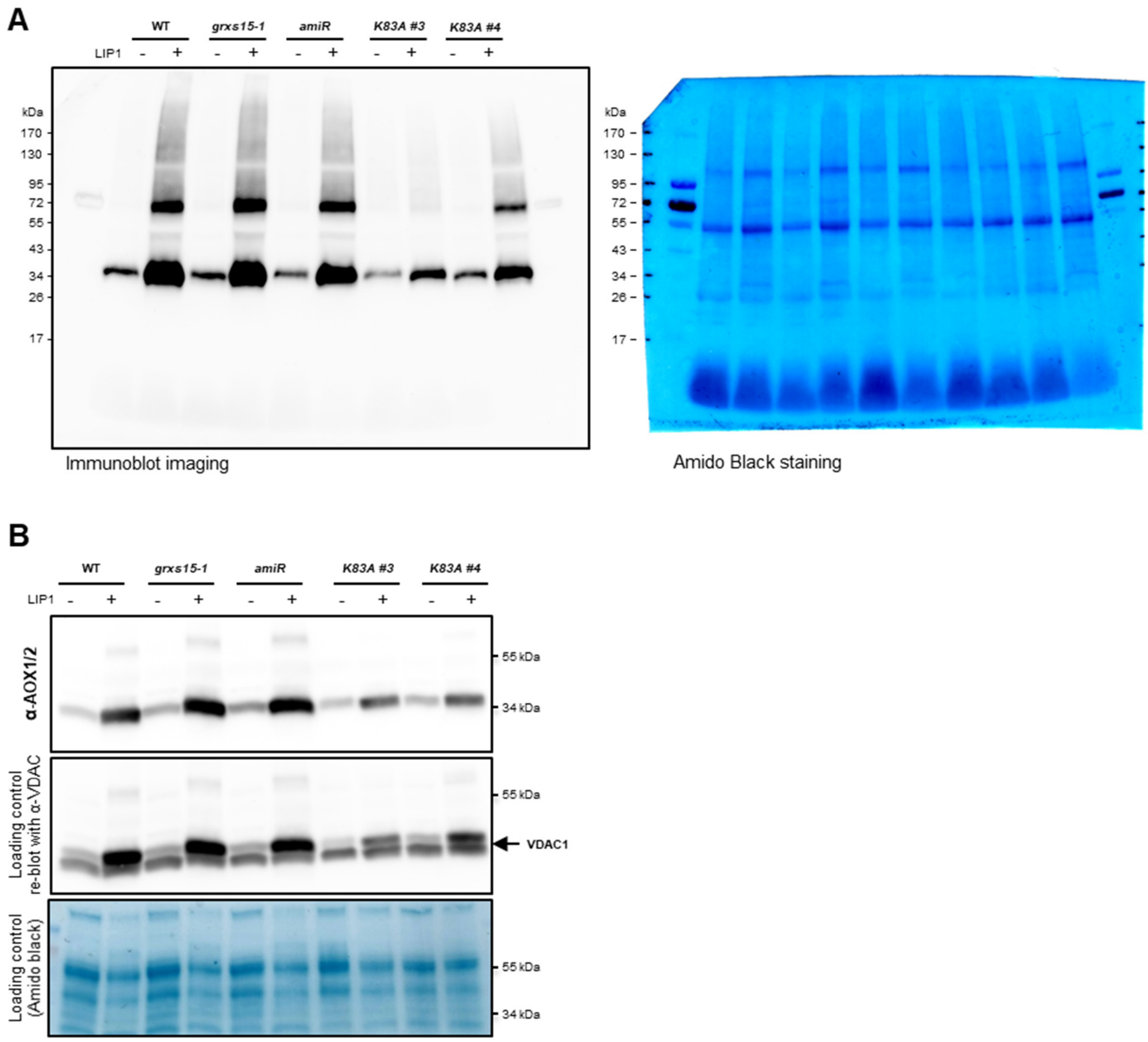


**Supplemental Figure S20. Overexpression of *LIP1* has an impact on the regulation of AOX expression. A,** Protein gel blot analysis with primary antibodies raised against AOX1/2 (AS04 054; Agrisera). 15 µg of proteins extracted from mitochondria isolated from leaves of 7-week-old soil-grown plants. Amido Black stain serves as loading control. **B,** Similar to (**A**), protein gel blot analysis with primary antibodies raised against AOX1/2 with 15 µg of protein extracted from mitochondria isolated from seedlings grown in hydroponic culture for 16 days. Re-blot with antibody raised against VDAC1 and Amido black stain serves as loading control.

**Supplemental Table S1. Primers for genotyping.**

| Line name and ID | Allele type | Primer ID | Sequence (5’ → 3’) |
|---|---|---|---|
| ***grxs15-1*** | WT | #2710 | GGAGATTCAGGGACACCTTTC |
| (SALK_112767) | | #2711 | ATGGTCCACTTCGTATGTTGG |
| | T-DNA | #1401 | ATTTTGCCGATTTCGGAAC |
| | | #2711 | ATGGTCCACTTCGTATGTTGG |
| ***amiR*** (Ströher et al., 2016) | T-DNA | #3791 | CGGCAACAGGATTCAATCTTAAG |
| | | #4428 | CGCACAATCCCACTATCCTT |
| ***grxs15-3*** | WT | #2708 | TGAAGCATACTTTTGGGATGG |
| (GK-837C05) | | #2709 | ATTCAAAACCATACGCTCACG |
| | T-DNA | #2709 | ATTCAAAACCATACGCTCACG |
| | | #432 | CCCATTTGGACGTGAATGTAGACAC |
| ***K83A T-DNA*** (Moseler et al., 2015) | T-DNA | #2842 | AGGGACACCAGCCATGTAGAT |
| | | #3613 | GTTTTCCCAGTCACGACGTTGT |
| ***aco3*** | WT | #4974 | CACTGTCTCATCGCTTCTTCC |
| (SALK_014661) | | #4975 | TCCAACAAAATCAATCCCTTG |
| | T-DNA | #4975 | TCCAACAAAATCAATCCCTTG |
| | | #1401 | ATTTTGCCGATTTCGGAAC |
| ***+OAS-TL C*** | T-DNA | #5127 | CGATGATCATGGCTTCAAGG |
| (pMDC32_$35S_{pro}$:*OAS-TL C*) | | #3791 | CGGCAACAGGATTCAATCTTAAG |

**Supplemental Table S2. Primers used for cloning.**

| Gene | Purpose | Primer ID | Sequence (5’ → 3’) |
|---|---|---|---|
| *LIP1* (AT2G20860) | Cloning in pSS01 | #4874 | GGGGACAAGTTTGTACAAAAAAGCAGGC<br>TTAATGCATTCGCGCTCCGCC |
| | | #4877 | GGGGACCACTTTGTACAAGAAAGCTGGG<br>TGCGGGGATGTAGAAGGAGAAG |
| | Cloning in pB7WG2 | #4874 | GGGGACAAGTTTGTACAAAAAAGCAGGC<br>TTAATGCATTCGCGCTCCGCC |
| | | #4876 | GGGGACCACTTTGTACAAGAAAGCTGGG<br>TCTACGGGGATGTAGAAGGAGA |

Primer #4874 includes a start codon and part the DNA coding for the original signal peptide for mitochondrial targeting. Primer #4877 lacks a stop codon to allow translational fusion of LIP1 with a C-terminal GFP. Primer #4876 includes the stop codon.

**Supplemental Table S3. *grxs15* mutants overexpressing *LIP1-GFP* show a Mendelian segregation.**

| Genetic background | Line | *n* | BASTA-resistant | BASTA-sensitive | Resistant/total (%) | □$^{2}$ |
|---|---|---|---|---|---|---|
| ***WT+LIP1-GFP*** | **#2** | 235 | 176 | 59 | 74.9 | 0.001 |
| ***WT+LIP1-GFP*** | **#6** | 207 | 157 | 50 | 75.8 | 0.079 |
| ***WT+LIP1-GFP*** | **#8** | 105 | 83 | 22 | 79.0 | 0.917 |
| ***grxs15-1+LIP1-GFP*** | **#5** | 182 | 134 | 48 | 73.6 | 0.183 |
| ***grxs15-1+LIP1-GFP*** | **#6** | 155 | 116 | 39 | 74.8 | 0.002 |
| ***grxs15-1+LIP1-GFP*** | **#12** | 182 | 139 | 43 | 76.4 | 0.183 |
| ***amiR+LIP1-GFP*** | **#6** | 285 | 220 | 65 | 77.2 | 0.731 |
| ***amiR+LIP1-GFP*** | **#9** | 104 | 80 | 24 | 76.9 | 0.205 |
| ***amiR+LIP1-GFP*** | **#10** | 183 | 137 | 46 | 74.9 | 0.002 |
| ***K83A #3+LIP1-GFP*** | **#1** | 148 | 117 | 31 | 79.1 | 1.297 |
| ***K83A #3+LIP1-GFP*** | **#2** | 179 | 132 | 47 | 73.7 | 0.151 |
| ***K83A #3+LIP1-GFP*** | **#3** | 104 | 80 | 24 | 76.9 | 0.205 |
| ***K83A #4+LIP1-GFP*** | **#1** | 104 | 79 | 25 | 76.0 | 0.051 |
| ***K83A #4+LIP1-GFP*** | **#2** | 260 | 191 | 69 | 73.5 | 0.328 |
| ***K83A #4+LIP1-GFP*** | **#3** | 151 | 111 | 40 | 73.5 | 0.179 |

All three independent lines for overexpression of *LIP1-GFP* in *WT*, *grxs15-1*, *amiR*, *K83A #3,* and *#4* segregated with a ratio of about 3:1 for resistance of $T_2$ plants to BASTA after 10 days. □$^{2}$ was calculated on an expected segregation of 3:1 (α = 0.05).

**Supplemental Table S4 | Wild type lines overexpressing *LIP1* without GFP tag have a Mendelian segregation**

| Genetic background | Line | *n* | BASTA-resistant | BASTA-sensitive | Resistant/total (%) | □$^{2}$ |
|---|---|---|---|---|---|---|
| ***WT+LIP1*** | **#1** | 181 | 129 | 52 | 71.3 | 1.343 |
| ***WT+LIP1*** | **#2** | 168 | 133 | 35 | 79.2 | 1.556 |
| ***WT+LIP1*** | **#3** | 196 | 149 | 47 | 76.0 | 0.109 |

All three independent lines for overexpression of *LIP1* in WT segregated with a ratio of about 3:1 for resistance of $T_2$ plants to BASTA after 10 days. □$^{2}$ was calculated on an expected segregation of 3:1 (α = 0.05).

**Supplemental Table S5 | WT+*LIP1* lines overexpressing *OAS-TL C* show Mendelian segregation of *OAS-TL C***

| Genetic background | Line | *n* | T-DNA (*OAS-TL C*) positive | T-DNA negative | T-DNA positive/total (%) | □$^{2}$ |
|---|---|---|---|---|---|---|
| **WT+*LIP1+OAS-TL C*** | **#1** | 12 | 8 | 4 | 66.7 | 0.444 |
| **WT+*LIP1+OAS-TL C*** | **#2** | 24 | 18 | 6 | 75.0 | 0.000 |
| **WT+*LIP1+OAS-TL C*** | **#3** | 12 | 10 | 2 | 83.3 | 0.444 |

All three independent lines for overexpression of *OAS-TL C* in WT *and* WT+*LIP1* segregated with a ratio of about 3:1 for the presence of the *OAS-TL C* construct (primers #5127 and #2791, Supplementary Table 1). □$^{2}$ was calculated on an expected segregation of 3:1 (α = 0.05).

**Supplemental Table S6 | Primers used for quantitative assessment of expression levels by qRT-PCR**

| Gene | Primer ID | Sequence |
|---|---|---|
| ***LIP1*** | #5199 | CACCAGATGCCTTCGAGAGGTA |
| AT2G20860 | #5200 | TCTCCCGCTTTGTACGACGAC |
| ***ACO3*** | #5252 | ATGCTTGTTGTGCCTCCTGG |
| AT2G05710 | #5253 | TCTGGGTAGAGAAGGCCTTTGG |
| ***ACO2*** | #5254 | CAAGCTAGGAATTCCACTCCGGTT |
| AT4G26970 | #5255 | TCTCCACCACCAGGTTTAGGAAGA |
| ***GRXS15*** | #5256 | CGCTGTGAAATCCTTCAGCCAC |
| AT3G15660 | #5257 | GCTCCAATTCACCTTCCTTGTGC |
| ***TIP41*** | #3892 | AATGCGTTTGACGCACTAGC |
| AT4G34270 | #3893 | GAGACGGCTTGCTCCTGAAT |
| ***SAND*** | #2895 | CCATATTGCAAGAAGTTTGCGCGTCTG |
| AT2G28390 | #2896 | GCAAGTCATCGGGATGGAGAGACG |
